# The Drosophila Baramicin polypeptide gene protects against fungal infection

**DOI:** 10.1101/2020.11.23.394148

**Authors:** M.A. Hanson, L.B. Cohen, A. Marra, I. Iatsenko, S.A. Wasserman, B. Lemaitre

## Abstract

The fruit fly *Drososphila melanogaster* combats microbial infection by producing a battery of effector peptides that are secreted into the haemolymph. Technical difficulties prevented the investigation of these short effector genes until the recent advent of the CRISPR/CAS era. As a consequence, many putative immune effectors remain to be characterized and exactly how each of these effectors contributes to survival is not well characterized. Here we describe a novel *Drosophila* antifungal peptide gene that we name *Baramicin A*. We show that *BaraA* encodes a precursor protein cleaved into multiple peptides via furin cleavage sites. *BaraA* is strongly immune-induced in the fat body downstream of the Toll pathway, but also exhibits expression in the nervous system. Importantly, we show that flies lacking *BaraA* are viable but susceptible to the enomopathogenic fungus *Beauveria bassiana*. Consistent with *BaraA* being directly antimicrobial, overexpression of *BaraA* promotes resistance to fungi and the IM10-like peptides produced by *BaraA* synergistically inhibit growth of fungi in vitro when combined with a membrane-disrupting antifungal. Surprisingly, *BaraA* males but not females display an erect wing phenotype upon infection. Collectively, we identify a new antifungal immune effector downstream of Toll signalling, improving our knowledge of the *Drosophila* antimicrobial response.

## Introduction

The innate immune response provides the first line of defence against pathogenic infection. This reaction is usually divided into three stages: i) the recognition of pathogens through dedicated pattern recognition receptors, ii) the activation of conserved immune signalling pathways and iii) the production of immune effectors that target invading pathogens [1,2]. The study of invertebrate immune systems has led to key observations of broad relevance, such as the discovery of phagocytosis [3], antimicrobial peptides (AMPs) [4], and the implication of Toll receptors in metazoan immunity [5]. Elucidating immune mechanisms, genes, and signalling pathways has greatly benefited from investigations in the fruit fly *Drosophila melanogaster*, which boasts a large suite of molecular and genetic tools for manipulating the system. One of the best-characterized immune reactions of *Drosophila* is the systemic immune response. This reaction involves the fat body (an analog of the mammalian liver) producing immune effectors that are secreted into the haemolymph. In *Drosophila*, two NF-κB signalling pathways, the Toll and Imd pathways, regulate most inducible immune effectors: the Toll pathway is predominantly activated in response to infection by Gram-positive bacteria and fungi [5,6], while the immune-deficiency pathway (Imd) responds to the DAP-type peptidoglycan most commonly found in Gram-negative bacteria and a subset of Gram-positive bacteria [7]. These two signalling pathways regulate a transcriptional program that results in the massive synthesis and secretion of humoral effector peptides [6,8]. Accordingly, mutations affecting the Toll and Imd pathways cause extreme susceptibilities to systemic infection that reflect the important contribution of these pathways to host defence. The best-characterized immune effectors downstream of these pathways are antimicrobial peptides (AMPs). AMPS are small and often cationic peptides that disrupt the membranes of microbes, although some have more specific mechanisms [9]. Multiple AMP genes belonging to seven well-characterized families are induced upon systemic infection [10]. However transcriptomic analyses have revealed that the systemic immune response encompasses far more than just the canonical AMPs. Many uncharacterized genes encoding small secreted peptides are induced to high levels downstream of the Toll and Imd pathways, pointing to the role for these peptides as immune effectors [11]. In parallel, MALDI-TOF analyses of the haemolymph of infected flies revealed the induction of 24 peaks – mostly corresponding to uncharacterized peptides – that were named “IMs” for Immune-induced Molecules (IM1-IM24) [8]. Many of the genes that encode these components of the immune peptidic secretome have remained largely unexplored. This is mainly due to the fact that these IMs belong to large gene families of small genes that were until recently difficult to disrupt by mutagenesis.

The CRISPR/Cas9 gene editing approach now allows the necessary precision to delete small genes, singly or in groups, providing the opportunity to dissect effector peptide functions. In 2015 a family of 12 related IM-encoding genes, unified under the name *Bomanins*, were shown to function downstream of Toll. Importantly, a deletion removing 10 out of the 12 Bomanins revealed their potent contribution to defence against both Gram-positive bacteria and fungi [12]. While Bomanins contribute significantly to Toll-mediated defence, their molecular functions are still unknown and it is unclear if they are directly antimicrobial [13]. Two other IMs encoding IM4 and IM14 (renamed *Daisho1* and *Daisho2*, respectively) were shown to contribute downstream of Toll to resistance against specific fungi. Interestingly, Daisho peptides bind to fungal hyphae, suggesting direct antifungal activity [14]. Finally a systematic knock-out analysis of *Drosophila* AMPs revealed that they play an important role in defence against Gram-negative bacteria and some fungi, but surprisingly little against Gram-positive bacteria [15]. An unforeseen finding from these recent studies is the high degree of AMP-pathogen specificity: this is perhaps best illustrated by the specific requirement for *Diptericin*, but not other AMPs, in defence against *Providencia rettgeri* [15,16]. Collectively, these studies in *Drosophila* reveal that immune effectors can be broad or specific in mediating host-pathogen interactions. Understanding the logic of the *Drosophila* effector response will thus require a careful dissection of the remaining uncharacterized immune induced peptides.

Previous studies identified an uncharacterized Toll-regulated gene called *IMPPP/CG33470*, which we rename *“BaraA”* (see below), that encodes several IMs, indicating a role in the humoral response. Here, we have improved the annotation of IMs produced by *BaraA* to include: IM10, IM12 (and its sub-peptide IM6), IM13 (and its sub-peptides IM5 and IM8), IM22, and IM24. Using a *BaraA* reporter, we show that *BaraA* is not only immune-induced in the fat body, but also expressed in the head, and nervous system tissue including the eyes, and ocelli. Importantly, we show that flies lacking *BaraA* are viable but susceptible to specific infections, notably by the entomopathogenic fungus *Beauveria bassiana*. Consistent with this, the IM10-like peptides produced by *BaraA* inhibit fungal growth in vitro when combined with the antifungal Pimaricin. Surprisingly, *BaraA* deficient males also display a striking erect wing behaviour upon infection. Collectively, we identify a new antifungal immune effector downstream of Toll signalling, improving our knowledge of the *Drosophila* antimicrobial response.

## Results

### BaraA is regulated by the Toll pathway

Previous microarray studies from De Gregorio et al. [11] suggest that *BaraA* is primarily regulated by the Toll pathway, with a minor input from the Imd pathway (**Fig**. 1A). Consistent with this, we found several putative NF-κB binding sites upstream of the *BaraA* gene (guided by previous reports [17–19]). Notably there are two putative binding sites for Relish, the transcription factor of the Imd pathway and three putative binding sites for the Dif/Dorsal transcription factors acting downstream of Toll (**Fig**. S1A). We challenged wild type flies and Imd or Toll pathway mutants (*Rel^E20^* and *spz^rm7^* respectively) with the Gram-negative bacterium *Escherichia coli*, the yeast *Candida albicans*, or the Gram-positive bacterium *Micrococcus luteus*. RT-qPCR analysis confirms that *BaraA* is induced by infection with *E. coli, C. albicans*, or *M. luteus* (**Fig**. 1B and **Fig**. S1B). *BaraA* remains highly inducible in a *Relish* mutant background, albeit at slightly reduced level compared to the wild type. However *BaraA* expression is abolished in *spz^rm7^* flies. Collectively, the expression pattern of *BaraA* is reminiscent of the antifungal peptide gene *Drosomycin* with a primary input by the Toll pathway and a minor input from the Imd pathway [10,20].

**Figure 1:**
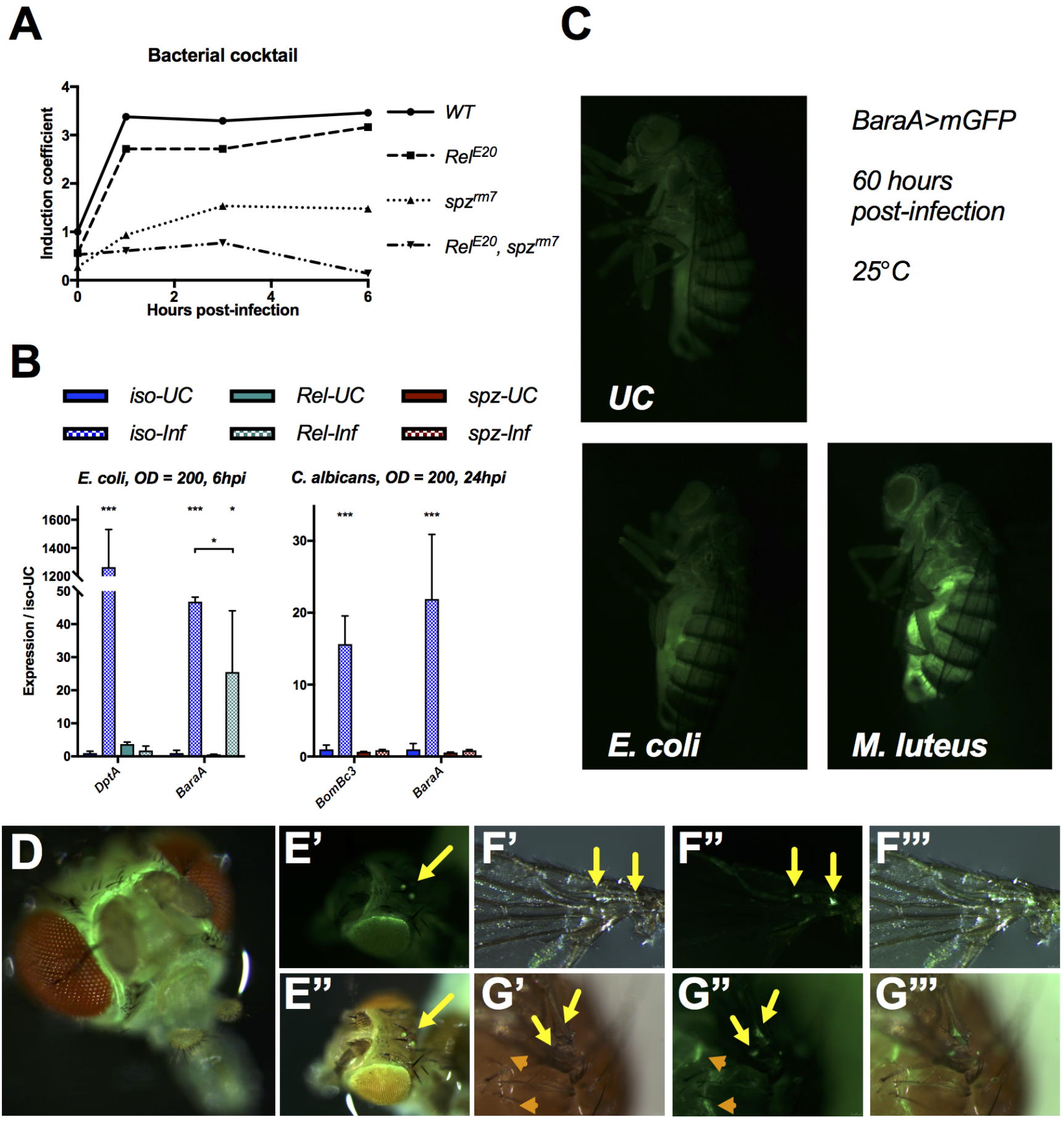
*BaraA* is an immune-induced gene regulated by the Toll pathway. A) Expression profile of *BaraA* upon bacterial challenge by a mixture of *E. coli* and *M. luteus* (from De Gregorio et al. [11]). Induction coefficient is a complex calculation of log-fold change reported in De Gregorio et al. [11], and values are given relative to wild type unchallenged expression levels. B) *BaraA* expression profiles in wild-type, *spz^rm7^* and *Rel^E20^* flies upon septic injury with the Gram-negative bacterium *E. coli* and the yeast *C. albicans. DptA* and *BomBc3* were used as inducible control genes for the Imd and Toll pathways respectively. Floating asterisks indicate significance relative to *iso-UC* where: * = p < .05 and, *** = p < .001. C) Use of a *BaraA* reporter reveals that *BaraA* induction upon infection is primarily driven by the fat body in adults, and results in a much stronger GFP signal upon pricking with *M. luteus* (which stimulates the Toll pathway) compared to *E. coli* (which stimulates the Imd pathway). Representative images taken 60h (adults) after handling alone (UC) or infection with *E. coli* or *M. luteus* (OD=200). D-G) *BaraA>mGFP* is highly expressed in the head (D), at the border of the eyes and in the ocelli (E), in the wing veins (F-G yellow arrows), and beneath the cuticle in the thorax (H, orange arrows).

To further characterize the expression of *BaraA*, we generated a *BaraA-Gal4* transgene in which 1675bp of the *BaraA* promoter sequence is fused to the yeast transcription factor Gal4. Use of *BaraA-Gal4>UAS-mCD8-GFP* (referred to as *BaraA>mGFP*) reveals that *BaraA* is strongly induced in the fat body 60h post infection by *M. luteus*, but less so by *E. coli* pricking (**Fig**. 1C); dissections confirmed this GFP signal is produced by the fat body. Larvae pricked with *M. luteus* also show a robust GFP signal primarily stemming from the fat body when examined 2hpi (**Fig**. S1C). We also observed a strong constitutive GFP signal in the headcase of adults (**Fig**. 1D), including the border of the eyes and the ocelli (**Fig**. 1E). Dissection confirmed that the *BaraA* reporter is expressed in brain tissue, notably in the central ventral brain furrow. Other consistent signals include GFP in the wing veins and subcutaneously along borders of thoracic pleura in adults (**Fig**. 1F–G), and in spermatheca of females (**Fig**. S1D). There was also sporadic GFP signal in other tissues that included the larval hindgut, the dorsal abdomen of developing pupae, and the seminal vesicle of males. These expression patterns largely agree with data reported in FlyAtlas1 [21].

### Baramicin A encodes a precursor protein cleaved into multiple peptides

Previous studies using bioinformatics and proteomics have suggested that four highly immune-induced peptides (IM10, IM12, IM13, and IM24) are encoded in tandem as a single polypeptide precursor by *IMPPP/BaraA* [8,22]. Some less-abundant sub-peptides (IM5, IM6, and IM8) are also produced by additional cleavage of IM12 and IM13 [22]. Using a newly generated null mutant (*“ΔBaraA,”* described below), we analyzed haemolymph samples of wild type and *ΔBara* flies infected with a bacterial mixture of *E. coli* and *M. luteus* by MALDI-TOF analysis. We confirmed the loss of the seven immune-induced peaks corresponding to IMs 5, 6, 8, 10, 12, 13, and 24 in *ΔBaraA* flies (**Fig**. 2A). We also noticed that an additional immune-induced peak at ~5975 Da was absent in our *BaraA* mutants. Upon re-visiting the original studies that annotated the *Drosophila* IMs, we realized this peak corresponded to IM22, whose sequence was never determined [8,22] (see supplementary information for details). We subjected haemolymph from infected flies to LC-MS proteomic analysis following trypsin digestion and found that in addition to the known IMs of *BaraA* (IMs 5, 6, 8, 10, 12, 13, and 24), trypsin-digested fragments of the *BaraA* C-terminus peptide were also detectable in the haemolymph (**Fig**. S2). The range of detected fragments did not match the full length of the C-terminus exactly, as the first four residues were absent in our LC-MS data (a truncation not predicted to arise via trypsin cleavage). The *BaraA* C-terminus lacking these four residues has a calculated mass of 5974.5 Da, exactly matching the observed mass of the IM22 peak absent in *BaraA* mutant flies. Furthermore in other *Drosophila* species these four residues of the *BaraA* C-terminus are instead an RXRR furin cleavage motif (**Fig**. S3A). Therefore IM22 cleavage in other species, even by an alternate cleavage process, should result in the same maturated IM22 domain as found in *D. melanogaster*. Taken together, we conclude that IM22 is the mature form of the BaraA protein C-terminus.

**Figure 2:**
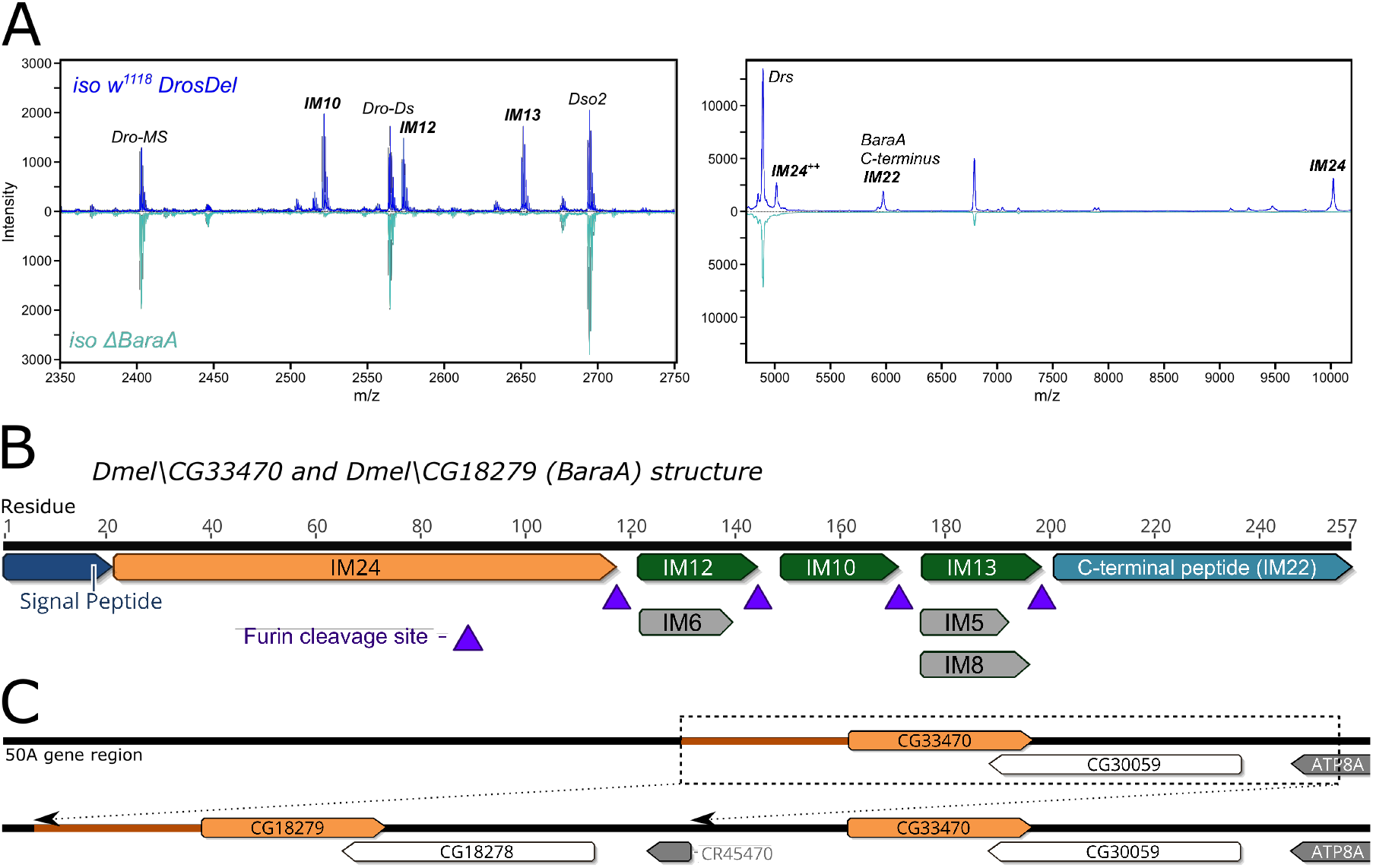
The *BaraA* gene structure. A) MALDI-TOF analysis of haemolymph from *iso w^1118^* wild-type and *iso ΔBaraA* flies 24 hours post-infection (hpi) confirms that *BaraA* mutants fail to produce the IM10-like and IM24 peptides. *iso ΔBaraA* flies also fail to produce an immune-induced peak at ~5794 Da corresponding to IM22 (the C-terminal peptide of BaraA, see supplementary information). B) The *BaraA* gene encodes a precursor protein that is cleaved into multiple mature peptides at RXRR furin cleavage sites. The sub-peptides IMs 5, 6, and 8 are additional minor cleavage products of IM12 and IM13. IM22 is additionally cleaved following its GIND motif (Fig. S2 and S3A). C) There is a *BaraA* locus duplication event present in the Dmel_R6 reference genome. This duplication is not fixed in laboratory stocks and wild-type flies (Hanson and Lemaitre, 2020; in prep.).

Thus, a single gene, *BaraA*, contributes to one third of the originally described *Drosophila* IMs. These peptides are encoded as a polypeptide precursor interspersed by furin cleavage sites (e.g. RXRR) (**Fig**. 2B). We note that the IM10, IM12 and IM13 peptides are tandem repeats of related peptides, which we collectively refer to as “IM10-like” peptides (**Fig**. S3B). The IM22 peptide also contains a similar motif as the IM10-like peptides (**Fig**. S3A-B), suggesting a related biological activity. We name this gene *“Baramicin A”* (symbol: *BaraA*) for the Japanese idiom Bara Bara 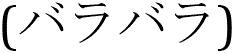, meaning “to break apart;” a reference to the fragmenting structure of the *Baramicin* precursor protein and its many peptidic products.

### A BaraA duplication is present in some laboratory stocks

Over the course of our investigation, we realized that *IMPPP (CG18279*) was identical to its neighbour gene *CG33470* owing to a duplication event of the *BaraA* locus present in the *D. melanogaster* reference genome. The exact nature of this duplication is discussed in a separate article (Hanson and Lemaitre; in prep). In brief, the duplication involves the entire *BaraA* gene including over 1kbp of 100% identical promoter sequence, and also the neighbouring sulfatase gene CG30059 and the 3’ terminus of the *ATP8A* gene region (**Fig**. 2C). We distinguish the two daughter genes as *BaraA1 (CG33470*) and *BaraA2 (CG18279*). Available sequence data suggests the *BaraA1* and *BaraA2* transcripts are 100% identical. We analyzed the presence of the *BaraA* duplication using a PCR assay spanning the junction of the duplicated region (supplementary data file 1). Interestingly, *BaraA* copy number is variable in common lab strains and wild flies, indicating this duplication event is not fixed in *D. melanogaster* (Hanson and Lemaitre; in prep).

### Over-expression of BaraA improves the resistance of immune deficient flies

*Imd, Toll* deficient flies are extremely susceptible to microbial infection as they fail to induce hundreds of immune genes, including antimicrobial peptides [11]. It has been shown that over-expression of even a single AMP can improve the resistance of *Imd, Toll* deficient flies [23]. As such, immune gene over-expression in *Imd, Toll* immune-compromised flies provides a direct assay to test the ability of a gene to contribute to defence independent of other immune effectors. We applied this strategy to *Baramicin A* by generating flies that constitutively express *BaraA* using the ubiquitous *Actin5C-Gal4* driver (*Act-Gal4*) in an immune-deficient *Rel^E20^, spz^rm7^* double mutant background (**Fig**. S4A). In these experiments, we pooled results from both males and females due to the very low availability of homozygous *Rel, spz* adults, particularly when combined with *Act-Gal4*. Overall, similar trends were seen in both sexes, and separate male and female survival curves are shown in **Fig**. S4.

Ubiquitous *BaraA* expression marginally improved the survival of *Rel, spz* flies upon infection with *M luteus* bacteria, however there was no effect upon infection with *E. coli* (**Fig. S4B-C**). On the other hand, ubiquitous expression of *BaraA* provided a more pronounced protective effect against infection by a variety of fungal pathogens. This was true upon pricking with *C. albicans* (**Fig**. 3A), or upon natural infections using *Aspergillus fumigatus* or *Neurospora crassa* filamentous fungi (**Fig**. 3B–C). This over-expression study reveals that *BaraA* alone can partially rescue the susceptibility of *Imd, Toll* deficient flies to infection, and points to a more prominent role for *BaraA* in antifungal defence.

**Figure 3:**
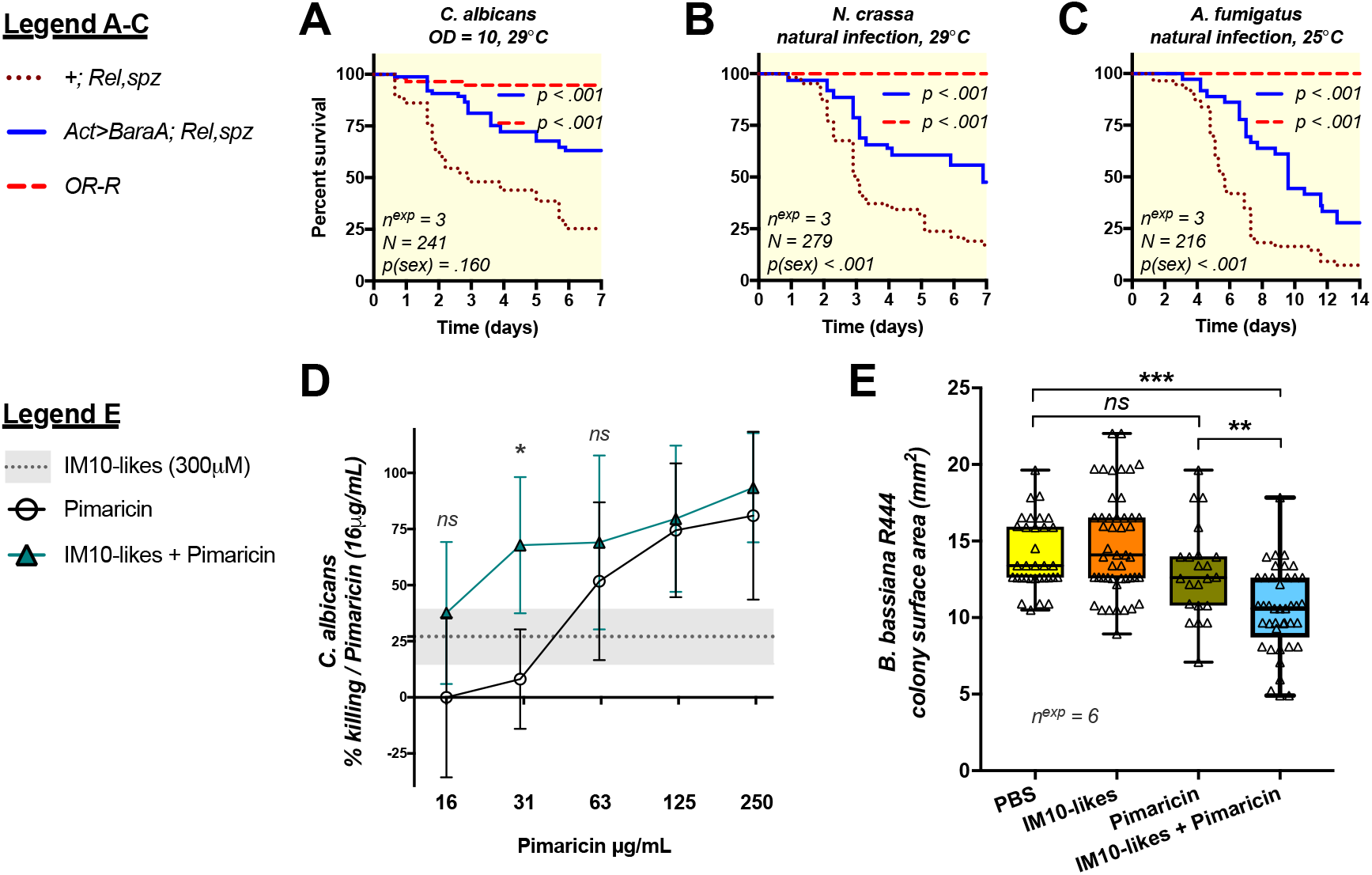
Overexpression of *BaraA* partially rescues the susceptibility of *Rel, spz* flies against fungi and BaraA IM10-like peptides inhibit fungal growth in vitro. A-C) Overexpression of *BaraA (Act>BaraA*) rescues the susceptibility of *Rel, spz* flies upon systemic infection with *C. albicans* (A), or natural infection with either *N. crassa* or *A. fumigatus* (B-C). Survivals represent pooled results from males and females (see Fig. S4 for sex-specific survival curves). D) A 300μM cocktail of the three IM10-like peptides improves the killing activity of the antifungal Pimaricin against *C. albicans* yeast. Error bars and the shaded area (IM10-likes alone) represent ±1 standard deviation from the mean. Killing activity (%) was compared against no-peptide controls, then normalized to the activity of Pimaricin alone. E) The IM10-like peptide cocktail also synergizes with Pimaricin to inhibit mycelial growth of *B. bassiana strain R444*. The diameters of individual colonies of *B. bassiana* were assessed after four days of growth at 25°C after peptide treatment, and surface area calculated as πr^2^.

### IM10-like peptides display antifungal activity in vitro

*The Baramicin A* gene encodes a polypeptide precursor that ultimately produces multiple mature peptides. However the most prominent *BaraA* products are the 23-residue IM10, 12, and 13 peptides (collectively the “IM10-like” peptides); indeed three IM10-like peptides are produced for every one IM24 peptide (**Fig**. 2B), and IM22 also bears an IM10-like motif (**Fig**. S3). This prompted us to explore the in vitro activity of the BaraA IM10-like peptides as potential AMPs.

We synthesized IM10, IM12, and IM13 and performed in vitro antimicrobial assays with these three IM10-like peptides using a 1:1:1 cocktail with a final concentration of 300μM (100 μM each of IM10, IM12, and IM13). Using a protocol adapted from Wiegand et al. [24], we monitored the microbicidal activity of this peptide cocktail either alone, or in combination with membrane-disrupting antibiotics that facilitate peptide entry into the cell. We based this approach on previous studies that showed that the microbicidal activities of Abaecin-like peptides, which target the bacterial DNA chaperone *DnaK*, increase exponentially in combination with a membrane disrupting agent [25–27]. We did not detect any killing activity of our IM10-like peptide cocktail alone against *Pectobacterium carotovora Ecc15* (hereafter *“Ecc15”), Enterococcus faecalis*, or *C. albicans*. We also found no activity of IM10-like peptides against *Ecc15 or E. faecalis* when co-incubated with a sub-lethal dose of Cecropin or Ampicillin respectively. However, we observed a synergistic interaction between IM10-like peptides and the antifungal Pimaricin against *C. albicans* (**Fig**. 3D). Co-incubation of the IM10-like cocktail with Pimaricin significantly improved the killing activity of Pimaricin at 32μg/mL relative to either treatment alone. While not statistically significant, the combination of IM10-like cocktail and Pimaricin also outperformed either the IM10-like cocktail alone or Pimaricin alone across the entire range of Pimaricin concentrations tested.

We next co-incubated dilute preparations of *B. bassiana* strain R444 spores under the same conditions as used previously with *C. albicans*, plated 2μL droplets, and assessed the diameters and corresponding surface area of colonies derived from individual spores after 4 days of growth at 25°C to assess growth rate. We found that neither the IM10-like cocktail nor Pimaricin alone affected surface area relative to PBS buffer control alone (Tukey’s HSD: p = 0.656 and 0.466 respectively). However in combination, the IM10-like cocktail plus Pimaricin led to significantly reduced colony size compared to either treatment alone, corresponding to a 19-29% reduction in surface area relative to controls (**Fig**. 3E, Tukey’s HSD: p < .01 in all cases). This indicates that incubation with IM10-like peptides and Pimaricin synergistically inhibits *B. bassiana* mycelial growth, revealing an otherwise cryptic antifungal effect of the BaraA IM10-like peptides.

Overall, we found that IM10-like peptides alone do not kill *C. albicans* yeast or impair *B. bassiana* mycelial growth in vitro. However, IM10-like peptides seem to synergize with the antifungal Pimaricin to inhibit growth of both of these fungi.

### BaraA deficient flies broadly resist like wild type upon bacterial infection

To further characterize *BaraA* function, we generated a null mutation of *BaraA* by replacing the ‘entire’ *BaraA* locus with a dsRed cassette using CRISPR mediated homology-directed repair with fly stocks that contain only one *BaraA* gene copy (BL2057 and BL51323). After isolation, this mutation (*BaraA^SW1^*) was then backcrossed once to a lab strain of *w^1118^* (used in [12–14]) to remove a second site mutation (see materials and methods). The resulting *w^1118^; BaraA^SW1^* flies are hereon referred to as *“w; ΔBaraA.”* Finally, the *BaraA^SW1^* mutation was isogenized by seven rounds of backcrossing into the *w^1118^ DrosDel* isogenic genetic background (*iso w^1118^*) [28] as described in Ferreira et al and are hereon referred to as “*iso ΔBaraA”* [29]. Relevant to this study, both our *OR-R* and *DrosDel iso w^1118^* wild type lines contain the duplication and thus have both *BaraA1 and A2* genes, while *w; ΔBaraA* and *iso ΔBaraA* flies lack *BaraA* entirely. In the following experiments, we compare the immune response of both *w; ΔBaraA* and *iso ΔBaraA* and focused only on phenotypes that were consistent in both genetic backgrounds.

We validated these mutant lines by PCR, qPCR and MALDI-TOF peptidomics (**Fig**. 2A, supplementary data file 1). *BaraA*-deficient flies were viable with no morphological defect. Furthermore, *ΔBaraA* flies have wild type Toll and Imd signalling responses following infection, indicating that *BaraA* is not required for the activation of these signaling cascades (**Fig**. S5A-C). Moreover, *BaraA* mutant flies survive clean injury like wild type (**Fig**. S5D), and have comparable lifespan to wild type flies (**Fig**. S5E). We next challenged *BaraA* mutant flies using our two genetic backgrounds with a variety of pathogens. We included susceptible Imd deficient *Rel^E20^* flies, Toll deficient *spz^rm7^* flies and *Bomanin* deficient *Bom^Δ55C^* flies as comparative controls. We observed that *BaraA* null flies have comparable resistance as wild type to infection with the Gram-negative bacterium *Ecc15* (**Fig**. S6A), or with the Gram-positive bacterium *B. subtilis* (**Fig**. S6B). In contrast, we saw a mild increase in the susceptibility of *w; ΔBaraA* flies to infection by the Gram-positive bacterium *E. faecalis* (HR = +0.73, p = .014). We also saw an early mortality phenotype in *iso ΔBaraA* flies (at 3.5 days, p < .001), although this was not ultimately statistically significant (**Fig**. 4A; p = .173). This mild susceptibility was also observed using flies carrying the *BaraA* mutation over a deficiency (*ΔBaraA/Df(BaraA)*), as well as in flies ubiquitously expressing *BaraA* RNAi (**Fig**. S7); however none of these sets of survival experiments individually reached statistical significance. Overall, the susceptibility of *BaraA* mutants to *E. faecalis* is mild, but consistent using a variety of genetic approaches.

**Figure 4:**
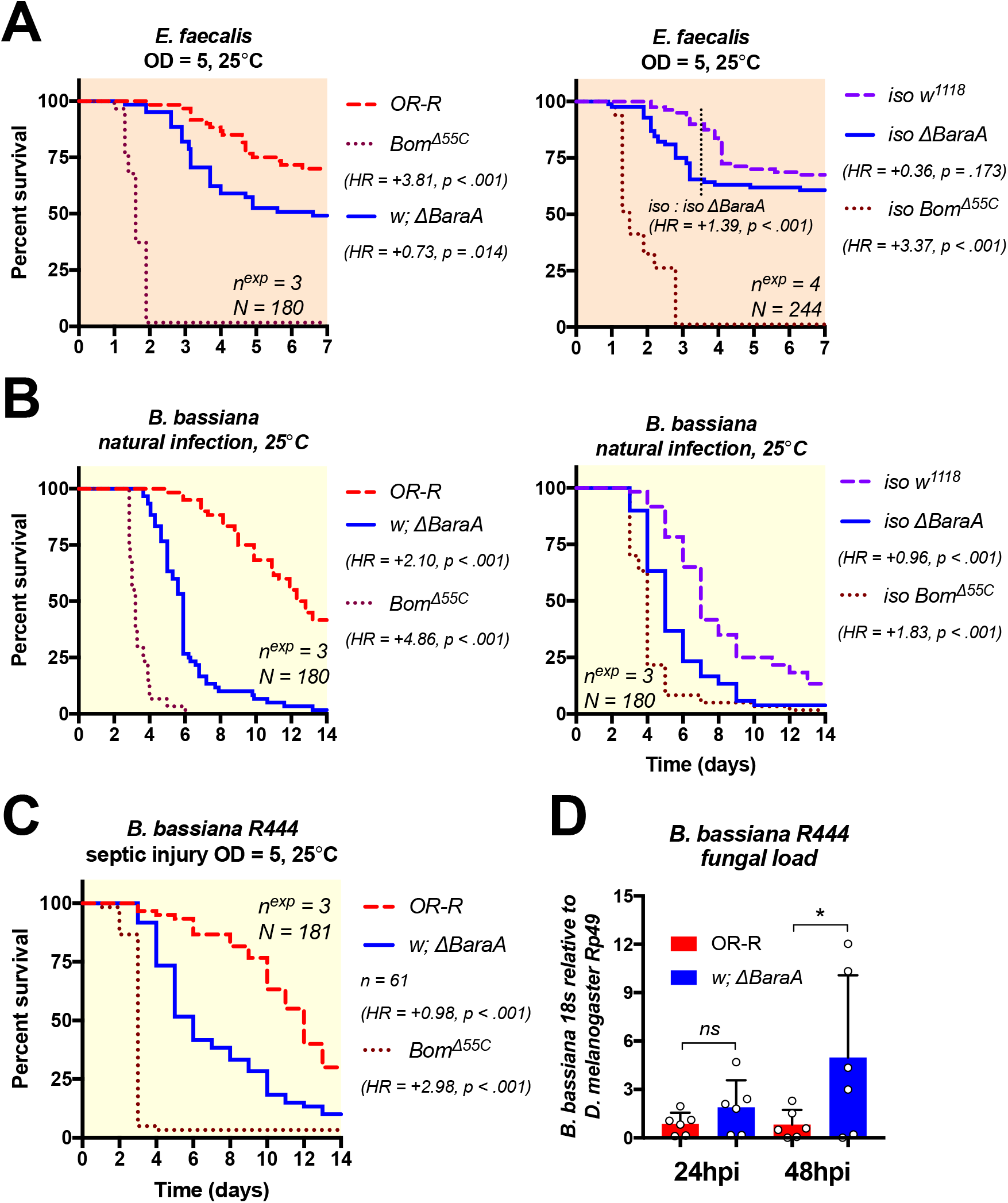
*ΔBaraA* flies are most susceptible to fungal infection. A) *BaraA* mutants with two different genetic backgrounds (here called *w* or *iso*) display a mild susceptibility to systemic infection with *E. faecalis*. This presents as an earlier mortality in the *iso* background (dotted line, p < .001). However, survival in *iso ΔBaraA* flies was not significantly different from wild type at seven days post-infection (p = .173). B) *BaraA* mutants in both backgrounds are susceptible to natural infection with the entomopathogenic fungus *B. bassiana*. C-D) Increased susceptibility (C) and fungal load (D) of *ΔBaraA* flies upon systemic infection by *B. bassiana strain R444*.

### BaraA mutant flies are highly susceptible to Beauveria fungal infection

Entomopathogenic fungi such as *B. bassiana* represent an important class of insect pathogens [6]. They have the ability to directly invade the body cavity by digesting and crossing through the insect cuticle. The Toll pathway is critical to survive fungal pathogens as it is directly responsible for the expression of *Bomanin, Daisho, Drosomycin* and *Metchnikowin* antifungal effectors [12,14,15,30,31]. The fact that i) *BaraA* is Toll-regulated, ii) BaraA IM10-like peptides display antifungal activity in vitro, and iii) *BaraA* overexpression improves the resistance of *Imd, Toll* deficient flies against fungi all point to a role for *BaraA* against fungal pathogens.

We infected *BaraA* mutants and wild type flies by rolling flies in sporulating *B. bassiana* petri dishes. Strikingly, both *w; ΔBaraA* and *iso ΔBaraA* flies displayed a pronounced susceptibility to natural infection with *B. bassiana* (HR = +2.10 or +0.96 respectively, p < .001 for both) (**Fig**. 4B). An increased susceptibility to fungi was also observed using flies carrying the *BaraA* mutation over a deficiency (**Fig**. S8A) or that ubiquitously express *BaraA* RNAi (**Fig**. S8B). Moreover, constitutive *BaraA* expression (*Act-Gal4>UAS-BaraA*) in an otherwise wild type background improves survival to *B. bassiana* relative to *Act-Gal4>OR-R* controls (HR = −0.52, p = .010) (**Fig**. S8C). Finally, we used a preparation of commercial *B. bassiana R444* spores (BB-PROTEC, Andermatt Biocontrol) to perform controlled systemic infections by pricking flies with a needle dipped in spore solution. In these experiments we monitored both survival and fungal load using qPCR primers specific to the *B. bassiana* 18S rRNA gene [32]. As seen with natural infection, *BaraA* mutants were highly susceptible to *Beauveria* systemic infection (**Fig**. 4C). Moreover, *BaraA* mutants suffered increased fungal load by 48 hours after infection (**Fig**. 4D).

Collectively, our survival analyses point to an important role for *BaraA* in defence against the entomopathogenic fungus *B. bassiana*. Consistent with a direct effect of *BaraA* on fungi, we observe a susceptibility of *BaraA* mutants to infection that is correlated with increased proliferation of *B. bassiana*.

### BaraA contributes to antifungal defence independent of Bomanins

Use of compound mutants carrying multiple mutations in effector genes has shown that some of them additively contribute to host resistance to infection [15]. Compound deletions of immune genes can also reveal contributions of immune effectors that are not detectable via single mutant analysis [15,33,34]. Recent studies have indicated that *Bomanins* play a major role in defence against fungi [12,13], though their mechanism of action is unknown. It is possible that *Bomanin* activity relies on the presence of *BaraA*, or vice versa. This prompted us to investigate the interaction of *Bomanins* and *BaraA* in defence against fungi. To do this, we recombined the *Bom^Δ55C^* mutation (that removes a cluster of 10 *Bomanin* genes) with *ΔBaraA*. While natural infection with *Aspergillus fumigatus* did not induce significant mortality in *BaraA* single mutants (**Fig**. S6C-D), we observed that combining *ΔBaraA* and *Bom^Δ55C^* mutations increases fly susceptibility to this pathogen relative to *Bom^Δ55C^* alone (HR = −0.46, p = .003; **Fig**. 5A). We next exposed these *ΔBaraA, Bom^Δ55C^*, double mutant flies to natural infection with 30mg of commercial spores of *B. bassiana R444*, which we found to be a less virulent *Beauveria* infection model. This is equivalent to approximately 60 million spores, much of which are removed afterwards upon fly grooming. When using this infection method, we found that *BaraA* mutation markedly increases the susceptibility of *Bom^Δ55C^* mutant flies (HR = −0.89, p < .001), approaching *spz^rm7^* susceptibility (**Fig**. 5B).

**Figure 5:**
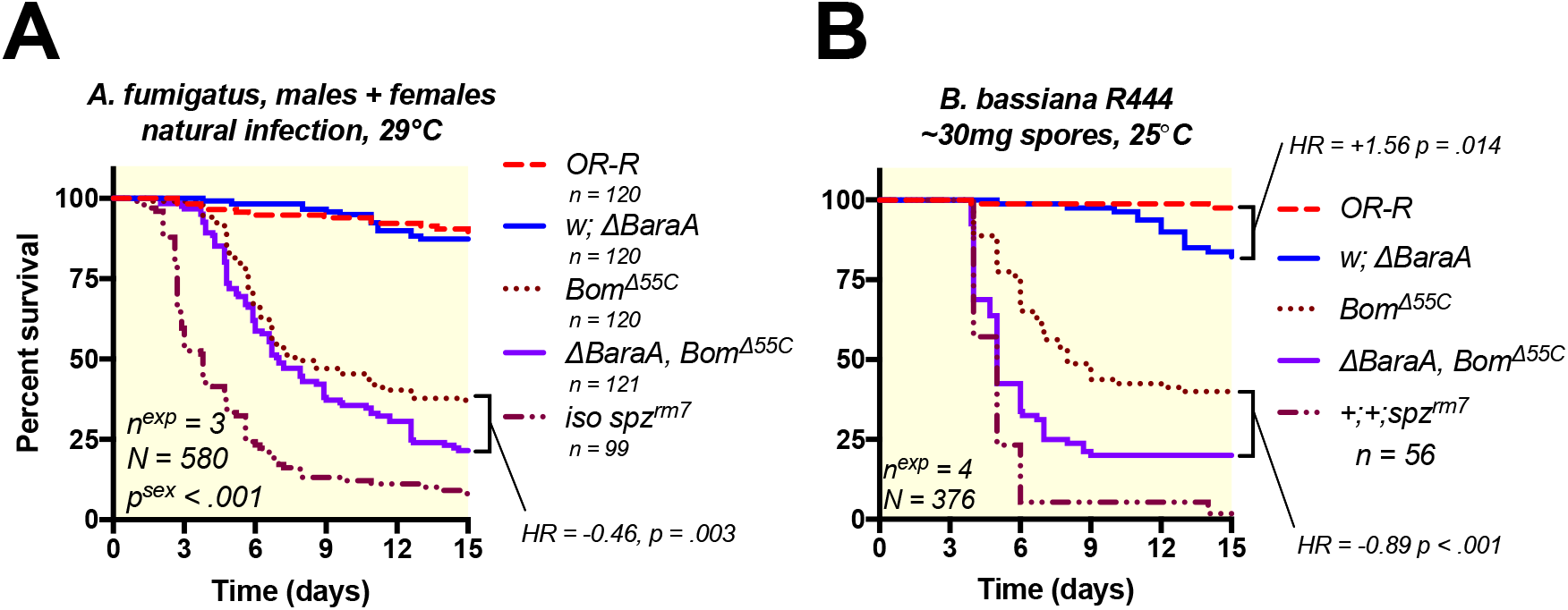
*BaraA* contributes to antifungal defence independent of *Bomanins*. A) *ΔBaraA, Bom^Δ55C^* double mutant flies were more susceptible than either mutation alone to natural infection with *A. fumigatus* (see Fig. S6C-D for sex-specific survival curves). B) *ΔBaraA, Bom^Δ55C^* double mutant flies were similarly more susceptible than individual mutants when given a mild (30mg of spores) *Beauveria* natural infection by *B. bassiana R444*.

We conclude that the contribution of *BaraA* to defence does not rely on the presence of *Bomanins*, and vice versa. This finding is consistent with the ability of constitutively expressed *BaraA* to improve survival outcome even in *Imd, Toll* deficient flies. Taken together, these results suggest *BaraA* improves survival against fungi independent of other effectors of the systemic immune response, consistent with a direct effect on invading fungi.

### ΔBaraA males display an erect wing phenotype upon infection

While performing infections with *A. fumigatus*, we observed a high prevalence of *BaraA* mutant flies with upright wings (**Fig**. 6A–B), a phenotype similar to the effect of disrupting the gene encoding the *“erect wing” (ewg*) transcription factor [35]. Curiously, this erect wing phenotype was most specifically observed in males. Upon further observation, erect wing was observed not only upon *A. fumigatus* infection, but also upon infections with all Gram-positive bacteria and fungi tested (**Table** S1). Increased prevalence of erect wing flies was observed upon infection with both live (**Fig**. 6C) and heat-killed *E. faecalis* (**Fig**. 6D), but less so upon clean injury or via infection with the Gram-negative bacteria *Ecc15* (**Fig**. S9A-B). Thus, the erect wing phenotype appears to be observed in *BaraA* mutants in response to stimuli known to activate the Toll pathway, but does not require a live infection.

**Figure 6:**
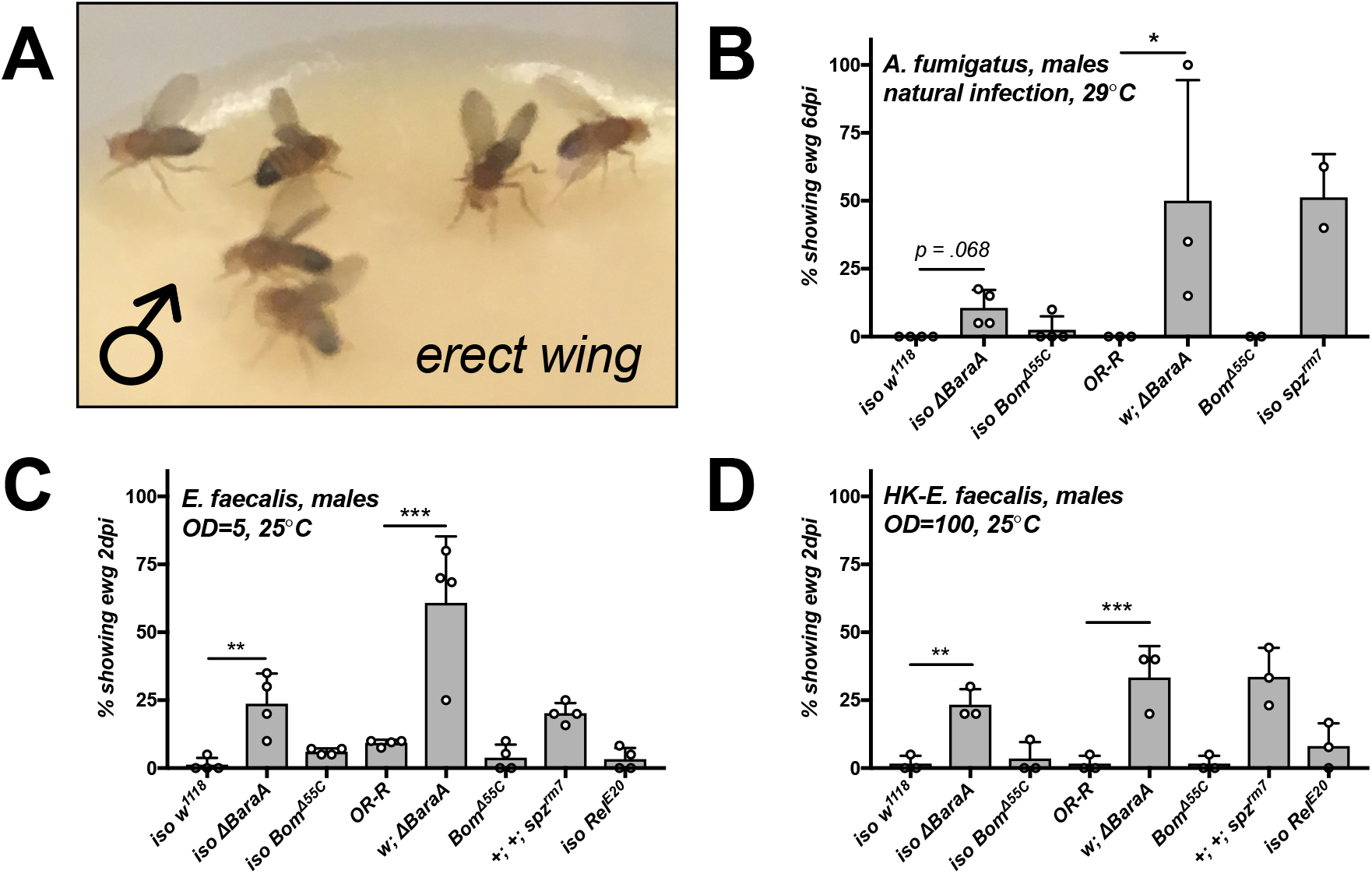
*ΔBaraA* males display an erect wing phenotype upon infection. A) *ΔBaraA* males displaying erect wing six days after *A. fumigatus* natural infection. B-D) *spz^rm7^* and *ΔBaraA* males, but not *Bom^Δ55C^* or *Rel^E20^* flies display the erect wing phenotype upon natural infection with *A. fumigatus* (B), or septic injury with live (C) or heat-killed *E. faecalis* (D). Barplots show the percentage of flies displaying erect wing following treatment, with individual data points reflecting replicate experiments. Additional challenges are shown in Table S1.

Such a phenotype in infected males has never been reported, but is reminiscent of the wing extension behaviour of flies infected by the brain-controlling “zombie” fungus *Enthomopthera muscae* [36]. Intrigued by this phenotype, we further explored its prevalence in other genetic backgrounds. We next confirmed that this phenotype was also observed in other *BaraA*-deficient backgrounds such as *Df(BaraA)/ΔBara;* however the penetrance was variable from one background to another. Erect wing was also observed in *ΔBaraA/+* heterozygous flies (*Df(BaraA)/+* or *ΔBaraA/+*), indicating that the lack of *BaraA* on one chromosome was sufficient to cause the phenotype (**Fig**. S9C and **Table** S1). Moreover, *spz^rm7^* flies that lack functional Toll signalling phenocopy *ΔBaraA* flies and display erect wing, but other immune-deficient genotypes such as mutants for the Toll-regulated *Bomanin* effectors (*Bom^Δ55C^*), or *Rel^E20^* mutants that lack Imd signalling, did not readily display erect wing (**Fig**. 6C–D, **Table** S1). Thus the erect wing phenotype is not linked to susceptibility to infection, but rather to loss of *BaraA* upon stimuli triggering the Toll immune pathway. This phenotype suggests an additional effect of *BaraA* on tissues related to the wing muscle or in the nervous system.

## Discussion

Seven *Drosophila* AMP families were identified in the 1980s-1990s either by homology with AMPs characterized in other insects or owing to their abundant production and microbicidal activities in vitro [37]. In the 2000s, genome annotations revealed the existence of many additional paralogous genes from the seven well-defined families of AMPs [38,39]. At that time, microarray and MALDI-TOF analyses also revealed the existence of many more small immune-induced peptides, which may function as AMPs [8,22]. Genetic analyses using loss of function mutations have recently shown that some of these peptides do play an important role in host defence, however key points surrounding their direct microbicidal activities remain unclear. In 2015, *Bomanins* were shown to be critical to host defence using genetic approaches, but to date no activity in vitro has been found [12,13]. Two candidate AMPs, Listericin [40] and GNBP-like3 [41], were shown to inhibit microbial growth upon heterologous expression using S2 cell lines or bacteria respectively. Most recently, Daisho peptides were shown to bind to fungal hyphae ex vivo, and are required for resisting fungal infection in vivo [14]. However the mechanism and direct microbicidal activity of these various peptides at physiological concentrations was not assessed.

In this study, we provide evidence from four separate experimental approaches that support adding *BaraA* to the list of bona-fide antifungal peptides. First, the *BaraA* gene is strongly induced in the fat body upon infection resulting in abundant peptide production. *BaraA* is also tightly regulated by the Toll pathway, which orchestrates the antifungal response. Second, loss of function study shows that *BaraA* contributes to resistance against fungi. *BaraA* mutation increases susceptibility to *B. bassiana*, and this is coupled with increased *B. bassiana* proliferation. Third, the antifungal activity of *BaraA* is independent of other key effectors. Over-expression of *BaraA* in the absence of other inducible peptides increased resistance of *Imd, Toll* deficient flies to various fungi including *C. albicans, A. fumigatus*, and *N. crassa*. Moreover *BaraA, Bomanin* double mutants suffered greater susceptibility than *Bomanin* mutants alone upon natural infection, even with a relatively avirulent fungal pathogen (*A. fumigatus*). Fourth, and lastly, a cocktail of the *BaraA* IM10-like peptides possesses antifungal activity against *C. albicans* and *B. bassiana* in vitro when co-incubated with the membrane disrupting antifungal Pimaricin.

While it is difficult to estimate the concentration of BaraA peptides in the haemolymph of infected flies, it is expected based on MALDI-TOF peak intensities that the IM10-like peptides should reach concentrations similar to other AMPs (up to 100μM) [10,20]; our in vitro assays used a peptide cocktail at the upper limit of this range. AMPs are often – but not exclusively – positively charged. This positive charge is thought to recruit these molecules to negatively charged membranes of microbes [10]. However the net charges at pH=7 of the IM10-like peptides are: IM10 +1.1, IM12 +0.1, and IM13 −0.9. Given this range of net charge, IM10-like peptides are not overtly cationic. However some AMPs are antimicrobial without being positively charged, exemplified by human Dermicidin [42] and anionic peptides of Lepidoptera that also synergize with membrane-disrupting agents [43]. More extensive in vitro experiments with additional fungi should confirm the range of BaraA peptide activities, and assay the potential activities of IM22 and IM24, which were not included in this study.

Our study also reveals that the *Baramicin A* gene alone produces at least 1/3 of the initially reported IMs. In addition to the IM10-like peptides and IM24 that were previously assigned to *BaraA* [22], we show IM22 is encoded by the C terminus of *BaraA*, and is conserved in other *Drosophila* species. The production of multiple IMs encoded as tandem repeats between furin cleavage sites is built-in to the BaraA protein design akin to a “protein operon.” Such tandem repeat organization is rare, but not totally unique among AMPs. This structure was first described in the bumblebee AMP Apidaecin [44], and has since also been found in Drosocin of *Drosophila neotestacea* [45]. In *D. melanogaster*, several AMPs are furin-processed including Attacin C and its pro-peptide MPAC, wherein both parts synergize in killing bacteria [25]. Therefore, furin cleavage in Attacin C enables the precise co-expression of distinct peptides with synergistic activity. It is interesting to note that IM10-like peptides did not show antifungal activity in the absence of membrane disruption by Pimaricin. An attractive hypothesis is that longer peptides encoded by BaraA such as IM22 and IM24 could contribute to the antifungal activity of *BaraA* by membrane permeabilization, allowing the internalization of IM10-like peptides. Indeed, the BaraA IM24 peptide is a short Glycine-rich peptide (96 AA) that is positively-charged (charge +2.4 at pH=7). These traits are shared by amphipathic membrane-disrupting AMPs such as Attacins [10], however the precise role for the Baramicin IM24 domain remains to be determined.

An unexpected observation of our study is the display of an erect wing phenotype by *BaraA* deficient males upon infection. Our study suggests that this is not a consequence of the genetic background, but rather relies on the activation of the Toll pathway in the absence of *BaraA*. Erect wing is also induced by heat-killed bacteria, and is not observed in *Bomanin* or *Relish* mutants, indicating that the erect wing phenotype is not a generic consequence of susceptibility to infection. The *erect wing* gene, whose inactivation causes a similar phenotype, is a transcription factor that regulates synaptic growth in developing neuromuscular junctions [35]. This raises the intriguing hypothesis that immune processes downstream of the Toll ligand Spaetzle somehow affect wing neuromuscular junctions, and that *BaraA* modulates this activity. Another puzzling observation is the sexual dimorphism exhibited for this response. Male courtship and aggression displays involve similar wing extension behaviours. Koganezawa et al. [46] showed that males deficient for *Gustatory receptor 32a (Gr32a*) failed to unilaterally extend wings during courtship display. *Gr32a*-expressing cells extend into the subesophageal ganglion where they contact mAL, a male-specific set of interneurons involved in unilateral wing display [46]. One possible explanation for the male specific effects of BaraA could be that *BaraA* mediates this effect through interactions with such male-specific neurons. *BaraA* is highly produced in the fat body upon infection but also expressed in the nervous system (**Fig**. 1D–E and FlyAtlas1 [21]). Further studies should decipher whether the preventative effect of *BaraA* on the erect wing phenotype is cell autonomous or linked to BaraA peptides secreted into the haemolymph. Recent studies have highlighted how NF-κB signalling in the brain is activated by bacterial peptidoglycan [47], and that immune effectors expressed either by fat body surrounding the brain or from within brain tissue itself affect memory formation [41]. Moreover, an AMP of nematodes regulates aging-dependent neurodegeneration through binding to its G-protein coupled receptor, and this pathway is sufficient to trigger neurodegeneration following infection [48]. Thus immune-inducible AMPs can have striking interactions with neurological processes. As such, future studies characterizing the role of *BaraA* in the erect wing phenotype should provide insight on interactions between systemic immunity and host physiology more generally.

Here we describe a complex immune effector gene that produces multiple peptide products. *BaraA* encodes many of the most abundant immune effectors induced downstream of the Toll signalling pathway, and indeed *BaraA* promotes survival upon fungal infection. How each peptide contributes to the immune response and/or erect wing behaviour will be informative in understanding the range of effects immune effectors can have on host physiology. This work and others also clarifies how the cocktail of immune effectors produced upon infection acts specifically during innate host defence reactions.

## Materials and methods

### Fly genetics and sequence comparisons

Sequence files were collected from FlyBase [49] and recently-generated sequence data [45,50] and comparisons were made using Geneious R10. Putative NF-κB binding sites were annotated using the Relish motif “GGRDNNHHBS” described in Copley et al. [18] and a manually curated amalgam motif of “GGGHHNNDVH” derived from common Dif binding sites described previously [17,19]. Gene expression analyses were performed using primers described in supplementary data file 1, and further microarray validation for *BaraA* expression comes from De Gregorio et al. [11].

The *UAS-BaraA* and *BaraA-Gal4* constructs were generated using the TOPO pENTR entry vector and cloned into the pTW or pBPGUw Gateway^™^ vector systems respectively. The *BaraA-Gal4* promoter contains 1675bp upstream of *BaraA1* (but also *BaraA2*, sequence in supplementary information). The *BaraA-Gal4* construct was inserted into the VK33 attP docking site (BDSC line #24871). The *BaraA^SW1^ (ΔBaraA*) mutant was generated using CRISPR with two gRNAs and an HDR vector by cloning 5’ and 3’ region-homologous arms into the pHD-dsRed vector, and consequently *ΔBaraA* flies express dsRed in their eyes, ocelli, and abdomen. *ΔBaraA* was generated using the Bloomington stocks BL2057 and BL51323 as these backgrounds contain only one copy of the *BaraA* locus. The induction of the immune response in these flies was validated by qPCR and MALDI-TOF proteomics, wherein we discovered an aberrant *Dso2* locus in these preliminary *BaraA^SW1^* flies. We thus backcrossed the *BaraA^SW1^* mutation once with a standard *w^1118^* background (used in [12–14]) and screened for wild type *Dso2* before use in any survival experiments. Additionally, *ΔBaraA* was isogenized into the *DrosDel w^1118^* isogenic background for seven generations before use in isogenic fly experiments as described in Ferreira et al. [29].

A full description of fly stocks used for crosses and in experiments is provided in supplementary data file 1.

### Microbe culturing conditions

Bacteria and *C. albicans* yeast were grown to mid-log phase shaking at 200rpm in their respective growth media (Luria Bertani, Brain Heart Infusion, or Yeast extract-Peptone-Glycerol) and temperature conditions, and then pelleted by centrifugation to concentrate microbes. Resulting cultures were diluted to the desired optical density at 600nm (OD) for survival experiments, which is indicated in each figure. The following microbes were grown at 37°C: *Escherichia coli strain ll06* (LB), *Enterococcus faecalis* (BHI), and *Candida albicans* (YPG). The following microbes were grown at 29°C: *Erwinia carotovora carotovora (Ecc15*) (LB) and *Micrococcus luteus* (LB). For filamentous fungi and molds, *Aspergillus fumigatus* was grown at 37°C, and *Neurospora crassa* and *Beauveria bassiana* were grown at room temperature on Malt Agar in the dark until sporulation. *Beauveria bassiana strain R444* commercial spores were produced by Andermatt Biocontrol, product: BB-PROTEC.

### Survival experiments

Survival experiments were performed as previously described [15], with 20 flies per vial with 2-3 replicate experiments. 3-5 day old males were used in experiments unless otherwise specified. As *Rel, spz* double mutant flies and largely wild type backgrounds differ drastically in their immune competence, we selected pathogens, infection routes, and temperatures to provide infection models that could best reveal phenotypes in these disparate genetic backgrounds. For fungi natural infections, flies were flipped at the end of the first day to remove excess fungal spores from the vials. Otherwise, flies were flipped thrice weekly. Statistical analyses were performed using a Cox proportional hazards (CoxPH) model in R 3.6.3. We report the hazard ratio (HR) alongside p-values as a proxy for effect size in survival experiments. Throughout our analyses, we required p < .05 as evidence to report an effect as significant, but note interactions with |HR| near or above 0.5 as potentially important provided p-value approached .05, and tamp down importance of interactions that were significant, but have relatively minor effect size (|HR| less than 0.5) in our discussion of the data.

### Erect wing scoring

The erect wing phenotype was scored as the number of flies with splayed wings throughout a distinct majority of the period of observation (30s); if unclear, the vial was monitored an additional 30s. Here we define splayed wings as wings not at rest over the back, but did not require wings to be fully upright; on occasion wings were held splayed outward at ~45° relative to the dorsal view, and often slightly elevated relative to the resting state akin to male aggressive displays. Sometimes only one wing was extended, which occurred in both thoracic pricking and fungi natural infections; these flies were counted as having erect wing. In natural infections, the typical course of erect wing display developed in two fashions at early time points, either: i) flies beginning with wings slightly splayed but not fully upright, or ii) flies constantly flitting their wings outward and returning them to rest briefly, only to flit them outward again for extended periods of time. Shortly after infection, some flies were also observed wandering around with wings beating at a furious pace, which was not counted as erect wing. However at later time points erect wing flies settled more permanently on upright splayed wings. Erect wing measurements were taken daily following infection, and erect wing flies over total flies was converted to a percent. Data points in **Fig**. 6B–D represent % with erect wing in individual replicate experiments with ~20 flies per vial. Flies stuck in the vial, or where the wings had become sticky or mangled were not included in totals. **Table** S1 reports mean percentages across replicate experiments for all pathogens and genotypes where erect wing was monitored. Days post-infection reported in **Table** S1 were selected as the final day prior to major incidents of mortality. For *E. faecalis* live infections, *Bom^Δ55C^* and *spz*^rm7^ erect wing was taken at 1dpi due to major mortality events by 2dpi specifically in these lines.

Erect wing measurements were performed in parallel with survival experiments, which often introduced injury to the thorax below the wing possibly damaging flight muscle. It is unlikely that muscle damage explains differences in erect wing display. First: we noticed erect wing initially during natural infections with *A. fumigatus*, and observed erect wing upon *B. bassiana R444* and *Metarhizium rileyi PHPl705* natural infections (**Table** S1; *M. rileyi* = NOMU-PROTEC, Andermatt Biocontrol). Second: only 1 of 75 total *iso w^1118^* males displayed erect wing across 4 systemic infection experiments with *E. faecalis*. For comparison: 19 of 80 total *iso ΔBaraA* and 48 of 80 *w; ΔBaraA* flies displayed erect wing (**Table** S1). Future studies might be better served using an abdominal infection mode, which can have different infection dynamics [51]. However we find erect wing display to be robust upon either septic injury or natural infection modes.

### IM10-like peptide in vitro activity

The 23-residue Baramicin peptides were synthesized by GenicBio to a purity of >95%, verified by HPLC. An N-terminal pyroglutamate modification was included based on previous peptidomic descriptions of Baramicins IM10, IM12, and IM13 [52], which we also detected in our LC-MS data (**Fig**. S2). Peptides were dissolved in DMSO and diluted to a working stock of 1200μM in 0.6% DMSO; the final concentration for incubations was 300μM in 0.15% DMSO. For microbe-killing assays, microbes were allowed to grow to log-growth phase, at which point they were diluted to ~50cells/μL. Two μL of culture (~100 cells), and 1μL water or antibiotic was mixed with 1μL of a 1:1:1 cocktail of IM10, IM12, and IM13 peptides to a final concentration of 300μM total peptides; 1μL of water + DMSO (final concentration = 0.15% DMSO) was used as a negative control. Four μL microbe-peptide solutions were incubated for 24h at 4°C. Microbe-peptide cultures were then diluted to a final volume of 100μL and the entire solution was plated on LB agar or BiGGY agar plates. Colonies were counted manually. For combinatorial assays with bacteria, *C. albicans* yeast, and *B. bassiana R444* spores, peptide cocktails were combined with membrane disrupting antimicrobials effective against relevant pathogens beginning at: 10 μM Cecropin A (Sigma), 500μg/mL ampicillin, or 250μg/mL Pimaricin (commercially available as “Fungin,” InVivogen), serially diluted through to 0.1 μM, 0.5μg/mL, and 4μg/mL respectively.

*Beauveria bassiana R444* spores were prepared by dissolving ~30mg of spores in 10mL PBS, and then 4μL microbe-peptide solutions were prepared as described for *C. albicans* followed by incubation for 24h at 4°C; this spore density was optimal in our hands to produce distinct individual colonies. Then, 4μL PBS was added to each solution and 2μL droplets were plated on malt agar at 25°C. Colony diameters were measured 4 days after plating by manually analyzing colony diameters in InkScape v0.92. Experimental batches were included as co-variates in one-way ANOVA analysis. The initial dataset approached violating Shapiro-Wilk assumptions of normality (p = 0.061) implemented in R 3.6.3. We subsequently removed four colonies from the analysis, as these outliers were over two standard deviations lower than their respective mean (removed colonies: PBS 0.15cm, PBS 0.25cm, IM10-like+Pimaricin 0.21cm, and a second IM10-like+Pimaricin colony of 0.21cm); the resulting Shapiro-Wilk p-value = 0.294, and both QQ and residual plots suggested a normal distribution. Final killing activities and colony surface areas were compared by One-way ANOVA with Holm-Sidak multiple test correction (*C. albicans*) and Tukey’s honest significant difference multiple test correction (*B. bassiana R444*).

### Gene expression analyses

RNA was extracted using TRIzol according to manufacturer’s protocol. cDNA was reverse transcribed using Takara Reverse Transcriptase. qPCR was performed using PowerUP mastermix from Applied Biosystems at 60°C using primers listed in supplementary data file 1. Gene expression was quantified using the PFAFFL method [53] with *Rp49* as the reference gene. Statistical analysis was performed by one-way ANOVA with Holm-Sidak’s multiple test correction or student’s t-test. Error bars represent one standard deviation from the mean.

### Proteomic analyses

Raw haemolymph samples were collected from immune-challenged flies for MALDI-TOF proteomic analysis as described in [14,15]. MALDI-TOF proteomic signals were confirmed independently at facilities in both San Diego, USA and Lausanne, CH. In brief, haemolymph was collected by capillary and transferred to 0.1% TFA before addition to acetonitrile universal matrix. Representative spectra are shown. Peaks were identified via corresponding m/z values from previous studies [8,22]. Spectra were visualized using mMass, and figures were additionally prepared using Inkscape v0.92.

## Supporting information

Hanson_etal_2020_supplemetary_figures_and_data_file_1

## Author contributions

MAH planned experiments, performed bioinformatic analyses, infection experiments, and in vitro assays. BL supervised the project and MAH and BL wrote the manuscript. LC planned and generated the *BaraA* deletion and performed key descriptive experiments and observations. AM assisted with infection and in vitro assays. MH, II, and SAW generated and supplied critical fly stock reagents and provided constructive commentary.

## Acknowledgements

This research was supported by Sinergia grant CRSII5_186397 and Novartis Foundation 532114 awarded to Bruno Lemaitre, and by National Institute of Health (NIH) grant R01 GM050545 to Steven Wasserman. We thank Jean-Philippe Boquete for assistance with the generation of Gal4 and UAS constructs. We would also like to acknowledge the technical expertise provided by the proteomics and mass spectrometry facilities in both UCSD and EPFL, and specifically Adrien Schmid. The name *“Baramicin’’* was partly inspired by Eichero Oda’s character “Buggy,” a Bara-Bara superhuman. Finally, we further thank Huang et al. (*in preparation*) for their cooperation in publishing initial descriptions of the *BaraA* gene, and for stimulating discussion.

## Supplemental figure and table captions

**Figure S1: Supplemental *BaraA* expression patterns. A**) 400bp of upstream sequence from *BaraA* annotated with putative *Rel* or *Dif/dl* binding sites (included in supplemental data file 1). **B**) Expression of *BaraA in wild-type* and *spz^rm7^ flies* following injury with the Gram-positive bacteria *M. luteus* . **C**) The *BaraA>mGFP* reporter line shows a robust induction of GFP 2hpi upon pricking with *M. luteus* in larvae. **D**) Expression of *BaraA>mGFP* in the spermatheca of females (yellow arrow). Representative images shown.

**Figure S2: LCMS coverage of trypsin-digested and detected BaraA peptides aligned to the protein coding sequence**. Peptide fragments cover the whole precursor protein barring furin site-associated motifs. Additionally, two peptide fragments are absent: i) the first 4 residues of the C-terminus (“GIND,” not predicted *a priori*), and ii) the C-terminus peptide’s “RPDGR” motif, which is predicted as a degradation product of Trypsin cleavage and whose size is beyond the minimum range of detection. Without the GIND motif, the mass of the contiguous C-terminus is 5974.5 Da, matching the mass observed by MALDI-TOF for IM22 (**Fig**. 2A). The N-terminal Q residues of IM10, IM12, IM13, and IM24 are pyroglutamate-modified, as described previously [22]. The asparagine residues of IM10-like peptides are sometimes deamidated, likely as a consequence of our 0.1% TFA sample collection method as “NG” motifs are deamidated in acidic conditions [54].

**Figure S3: Alignments of BaraA peptide motifs. A**) Aligned IM22 peptides of *Drosophila Baramicin A-like* genes, with the IM10-like ‘VWKRPDGRTV’ motif noted. The GIND residues at the N-terminus are cleaved off in *Dmel\BaraA* by an unknown process, and this site is similarly cleaved at RXRR furin cleavage site in subgenus Drosophila flies. As a consequence, the mature IM22 peptide is predicted to be the same across species even when different cleavage mechanisms are utilized. **B**) Alignment of the three IM10-like peptides of *D. melanogaster BaraA* with the “VXRPXRTV” motif noted.

**Figure S4: *Over-expression of BaraA partially* rescues *Rel, spz* double mutant susceptibility to infection in both males and females. A**) validation of the *UAS-BaraA* construct in the *Rel, spz* background. **B**) Overexpressing *BaraA* did not improve the survival of *Rel, spz* flies upon *E. coli* infection. **C**) Overexpressing *BaraA* only marginally improves survival of *Rel, spz* females, but not males, upon *M. luteus* infections. Infections using a higher dose tended to kill 100% of *Rel, spz* flies regardless of sex or expression of *BaraA*, suggesting that if *BaraA* overexpression does affect susceptibility to *M. luteus*, this effect is possible within only a narrow window of *M. luteus* concentration. **D-F**) Overexpressing *BaraA* improves survival of *Rel, spz* male and female flies upon injury with *C. albicans* (**D**) or natural infection with *A. fumigatus* (**E**) and *N. crassa* (**F**). P-values are shown for each biological sex in an independent CoxPH model not including the other sex relative to *Rel, spz* as a reference.

**Figure S5**: RT-qPCR shows that the expression of *BomBc3* (**A**) *Drs* (**B**) and *DptA* (**C**) is wild-type 18hpi in *iso ΔBaraA* flies. **D**) *BaraA* mutants survive clean injury like wild-type flies. **E**) *iso ΔBaraA* flies have similar lifespan compared with the *iso w^1118^* wild-type (males + females, *iso vs. iso ΔBaraA*: HR = 0.26, p = .118).

**Figure S6: Additional survivals using *ΔBaraA* flies in two distinct genetic backgrounds upon infection by a diversity of microbes. A**) No significant susceptibility of *ΔBaraA* flies to *Ecc15* infection. **B**) *w; ΔBaraA but not iso ΔBaraA* flies exhibit a marginal susceptibility to *B. subtilis* (HR > 0.5, p = .099). **C-D**) *w; ΔBaraA* males were slightly susceptible to *A. fumigatus* natural infection (HR > 0.5, p = .078), but not females, nor isogenic flies. Additional infections using *ΔBaraA, Bom^Δ55C^* double mutant flies reveal that *BaraA* mutation increases the susceptibility of *Bom^Δ55C^* flies in both males and females (cumulative curves shown in **Fig**. 5A). Blue backgrounds = Gram-negative bacteria, orange backgrounds = Gram-positive bacteria, yellow backgrounds = fungi.

**Figure S7: Additional survival analyses reveal only a minor contribution of *BaraA* to defence against infection by *E. faecalis*. A**) Crosses with a genomic deficiency (*Df(BaraA)*) leads to increased susceptibility in both the *w* background and isogenic DrosDel background, with *Df(BaraA)/ΔBaraA* flies suffering the greatest mortality in either crossing scheme. Both deficiency crosses yielded an earlier susceptibility in *BaraA*-deficient flies (shown with dotted black lines), however neither experiment ultimately reached statistical significance. **B**) *BaraA* RNAi flies (*Act>BaraA-IR*) suffered greater mortality than *Act>OR-R* or *OR-R/BaraA-IR* controls, but this was not statistically significant at α = .05; p-values reported are comparisons to *Act>BaraA-IR* flies.

**Figure S8: Additional survival analyses reveal a consistent contribution of *BaraA* to defence against natural infection with *B. bassiana*. A**) Crossing with a genomic deficiency (*Df(BaraA)*) leads to increased susceptibility of *Df(BaraA)/ΔBaraA* flies for both the *w* background and isogenic DrosDel background relative to wild-type controls (p < .05). **B**) *Act>BaraA-IR* flies were more susceptible than the *OR-R* wild-type (p = .008) and *OR>BaraA-IR* (p = .004), although not significantly different from our *Act>OR-R* control (p = .266). **C**) Overexpressing *BaraA (Act>UAS-BaraA*) improved survival against *B. bassiana* relative to *Act>OR-R* controls (HR = −0.52, p = 0.010).

**Figure S9: Frequency of erect wing display following additional challenges. A-B**) Erect wing frequencies 2dpi after clean injury (**A**), or *Ecc15* septic injury (**B**). The erect wing frequencies of flies pricked by HK-*E. faecalis* (**Fig**. 6D) are included in brown in **A** and **B** to facilitate direct comparison with the frequency observed upon Toll pathway activation. **C**) The frequency of erect wing display is increased following *E. faecalis* septic injury in *ΔBaraA/+* or *Df(BaraA)/+* flies. Data points are pooled from *w; ΔBaraA* and *iso ΔBaraA* crosses after *E. faecalis* infections shown in **Fig**. S7A and data in **Table** S1.

**Table S1: Erect wing frequencies from various infection experiments**. *Following initial erect wing observations upon* A. fumigatus *natural infection, we scored erect wing frequency in all subsequent survival experiments. Data represent the mean % of males displaying erect wing ± one standard deviation. n exp = number of replicate experiments performed, and dpi ewg taken = days post-infection where erect wing data were recorded. We additionally performed natural infections with* Metarhizium rileyi *that generally did not cause significant mortality even in* ΔBaraA, Bom^Δ55C^ *double mutant males, but nevertheless induced erect wing specifically in* ΔBaraA *males and* spz^rm7^ controls. Bacterial infections were performed by septic injury, while fungal challenges were natural infections performed by rolling flies in spores.

## Supplementary information

### Identification of the BaraA C-terminus as IM22 from Uttenweiler-Joseph et al

In 1998, Uttenweiler-Joseph et al. [8] described 24 immune-induced molecules by MALDI-TOF and informed predictions suggested that *BaraA* could encodes several of them [22]. We generated a knock out mutant for the *BaraA* gene (*BaraA^SW1^*), which we validated by MALDI-TOF peptidomic analysis. Strikingly, we noticed an immune-induced peak at ~5981 Da in Linear mode collections that is absent in *ΔBaraA* flies (**Fig**. 2A); this mass closely resembled the 5984 Da estimated mass of IM22 from Uttenweiler-Joseph et al. [8], for which sequence was never determined. We took the Linear masses reported for then-unknown IMs from Uttenweiler-Joseph et al. [8] and post-hoc generated a standard curve with now-confirmed mass values from Levy et al. [22]. Our post-hoc standard curve corrects the mass of IM22 as found in Uttenweiler-Joseph et al. [8] to be 5973.5 Da. Using the same approach with our own linear data we find a mass of 5975.1 Da for our 5981 Da peak (supplementary data file 1). With LCMS proteomics, we confirmed that the *BaraA* C-terminus is cleaved to remove 4 N-terminal residues, which should produce a putative 5974.5 Da peptide (**Fig** S2). Together these observations indicate the *BaraA* C-terminus encodes the following 53-residue mature peptide, matching the estimated mass of IM22: ARVQGENFVA RDDQAGIWDN NVSVWKRPDG RTVTIDRNGH TIVSGRGRPA QHY.

The *BaraA* gene is therefore involved in the production of over one third of the classical *Drosophila* IMs from Uttenweiler-Joseph et al. [8], including: IM5, 6, 8, 10, 12, 13, 20 (doubly-charged IM24 [22]), 22, and 24.

### Sequence of the BaraA-Gal4 promoter construct

The following 1675bp sequence was cloned from the DrosDel isogenic background into the pBPGUw vector to drive a downstream Gal4 gene, and inserted into the VK33 attP docking site using BDSC line #24871:

*Dif/dorsal binding site* (**bold**); **GGGHHNNDVH**

*Rel binding site* (<u>underline</u>): <u>GGRDNNHHBS</u>

>iso_DrosDel_BaraA_promoter-Gal4

CTGCTACTCCTCTACACATTCGACTCCTTCGCCTTGCTGGCTGGGAAAAAATTTTGCATA

ATTTATGTGGGTGCCGCGCACACGGAGGTCCCGACGGATTCGAAGTATCCGAAGGATTCG

AAAGGAAAACAACGCACGAGCACCACGGCCAACTGATTTAAATGCAATTGCACTGAAGT

ATTTTGTTTGGCGAACGAAGCTGGATGAAATAGGGGGGTGTGGGGTTTTCTATTGAGAC

ATCTGCACGTGCAACCGGAAACATCCGAAGAGAACAGCACAGGCCGGGCTACGCCGGGCA

ATTTCTTTTCATTTGCCAAGGTGTTGAGTTGCACCAACATTCGACATCGACGTGGCCAGA

AGCCAACAAAAGCCAAGAGCCAAACCCCTTTTTGTGGTCACAAGTGTCGTCTATTTGTCG

TGGGCATCTTGGGCACCTTGGGCATCCTCGACATCCTTGCCATTTTGGTCTGGCCAAGAC

AAACAACCAGCAAATTTAGTGTATTTTGTGCATTTTTAAAATTGTCCAAATTTATGTGA

CACGCTGCGCCAATTGATCAGATTAAATAAACATGAGGCCAAGCGAATCGAATTTGGCTT

CACCAAGAAGACAATGCAGTCTGTATTCAAATGGGTGGGCGCATCCACCAAGCGGTGAAT

ACAGTGACCGCTCGCTATAATGGACGGTCAGGTGTTACTTTAACTTAAAAAAATATGTA

ACAAATCTTATCAAGTTTGAAATAGATTGAAATAGATTTGGTTATTGCATTCGAAAGAT

ATATATTAAATTCGAATATTCCAAGAAATTTCATGAGAATGTCACTTATGTCATGAGAT

TATATTAACGTACGAATAAACAATGTATTTTCCAAAATTAAAAATAAAATTTAATTTAA

TTACGCAGTACCTTTACACTATCAGTCGGAGGTAATAACTCATATAATTAGATTAGCATT

AGATTTTAAAGCGAAAAACACTTAAAAGCTGAAATTATTAGACAACACTCTTAAATTAG

TCGAGCTGATATATAGCCTCAAGTTTTGCTTAAATCCAAAGATAAAGGAATGCCTTCAA

AAATATATTTTGTTTTATACCAAGTGACAGCAGAGAATGGGGTTGCAATATCTTAAAAG

AGTTTCACTTAGCCAATATTTACTGCCATTGTTGGCCACCAAATAGTAGCAACCAGAGAC

TTCCAGGAATATATTCTCGTGTCAAATGCAATCCACTTTAAATGCAACTATCTGGCGGCT

AAGAAAACCCGACAGTTTGATTCAAGTCGACGAAACAATATAAGCACGTGCTAAATAAA

GAGACCTATGCAGTTAATACTCTTGTCATATTATAATATAATTTAGTGACATAAGTTGC

ATGGTATACGAGTACTGAACAAGTTATGGCAGCTTTTCCAAATAAGCGATCACATATTCC

GCGGGATGATGGGTGGATTTCTAGCATATGTGGATGCTTAATGGCTTATTGC

GCGGGATGATGGGTGGATTTCTAGCATATGTGGATGCTTAATGGCTTATTGCGGGTCAG

GGCGGCGCAATCTGTTCAGAAATTCCCGAACGCACACCCATTTCAGATCAGATTGTGAC

GTTTTGGGAAATTCTTGACGATCGGTGTAAACAAGCTCAGCAACCAGATTCGATGGCTA

TTTGCCGGCTATAAATACTAGAAACCATTCGATTGCACTCAGTTGAAGCTGGGCTCTGGA

ACAGATCACA

## Notes

### Competing Interest Statement

The authors have declared no competing interest.

### Summary of Updates

Manuscript revisions and additional data added

## References

1. Lemaitre B, Hoffmann J. The Host Defense of Drosophila Melanogaster. Annual Review of Immunology. 2007;25: 697–743. doi:10.1146/annurev.immunol.25.022106.141615

2. Kurz CL, Ewbank JJ. Caenorhabditis elegans: An emerging genetic model for the study of innate immunity. Nature Reviews Genetics. 2003;4: 380–390. doi:10.1038/nrg1067

3. Kaufmann SHE. Immunology’s foundation: the 100-year anniversary of the Nobel Prize to Paul Ehrlich and Elie Metchnikoff. Nat Immunol. 2008;9: 705–712. doi:10.1038/ni0708-705

4. Steiner H, Hultmark D, Engström \AA, Bennich H, Boman HG. Sequence and specificity of two antibacterial proteins involved in insect immunity. Nature. 1981;292: 246–248. doi:10.1038/292246a0

5. Lemaitre B, Nicolas E, Michaut L, Reichhart JM, Hoffmann JA. The dorsoventral regulatory gene cassette spatzle/Toll/Cactus controls the potent antifungal response in Drosophila adults. Cell. 1996;86: 973–983. doi:10.1016/S0092-8674(00)80172-5

6. Lemaitre B, Reichhart JM, Hoffmann JA. Drosophila host defense: differential induction of antimicrobial peptide genes after infection by various classes of microorganisms. Proceedings of the National Academy of Sciences of the United States of America. 1997;94: 14614–9. doi:10.1073/pnas.94.26.14614

7. Lemaitre B, Kromer-Metzger E, Michaut L, Nicolas E, Meister M, Georgel P, et al. A recessive mutation, immune deficiency (imd), defines two distinct control pathways in the Drosophila host defense. Proc Natl Acad Sci U S A. 1995;92: 9465–9469. doi:10.1073/pnas.92.21.9465

8. Uttenweiler-Joseph S, Moniatte M, Lagueux M, Van Dorsselaer a, Hoffmann J a, Bulet P. Differential display of peptides induced during the immune response of Drosophila: a matrix-assisted laser desorption ionization time-of-flight mass spectrometry study. Proceedings of the National Academy of Sciences of the United States of America. 1998;95: 11342–11347. doi:10.1073/pnas.95.19.11342

9. Lazzaro BP, Zasloff M, Rolff J. Antimicrobial peptides: Application informed by evolution. Science. 2020;368. doi:10.1126/science.aau5480

10. Hanson MA, Lemaitre B. New insights on Drosophila antimicrobial peptide function in host defense and beyond. Curr Opin Immunol. 2020;62: 22–30. doi:10.1016/j.coi.2019.11.008

11. De Gregorio E, Spellman PT, Tzou P, Rubin GM, Lemaitre B. The Toll and Imd pathways are the major regulators of the immune response in Drosophila. EMBO Journal. 2002;21: 2568–2579. doi:10.1093/emboj/21.11.2568

12. Clemmons AW, Lindsay SA, Wasserman SA. An Effector Peptide Family Required for Drosophila Toll-Mediated Immunity. PLoS Pathogens. 2015;11. doi:10.1371/journal.ppat.1004876

13. Lindsay SA, Lin SJH, Wasserman SA. Short-Form Bomanins Mediate Humoral Immunity in Drosophila. J Innate Immun. 2018;10: 306–314. doi:10.1159/000489831

14. Cohen LB, Lindsay SA, Xu Y, Lin SJH, Wasserman SA. The Daisho Peptides Mediate Drosophila Defense Against a Subset of Filamentous Fungi. Front Immunol. 2020;11: 9. doi:10.3389/fimmu.2020.00009

15. Hanson MA, Dostálová A, Ceroni C, Poidevin M, Kondo S, Lemaitre B. Synergy and remarkable specificity of antimicrobial peptides in vivo using a systematic knockout approach. eLife. 2019;8. doi:10.7554/elife.44341

16. Unckless RL, Howick VM, Lazzaro BP. Convergent Balancing Selection on an Antimicrobial Peptide in Drosophila. Current Biology. 2016;26: 257–262. doi:10.1016/j.cub.2015.11.063

17. Busse MS, Arnold CP, Towb P, Katrivesis J, Wasserman SA. A κB sequence code for pathway-specific innate immune responses. EMBO Journal. 2007;26: 3826–3835. doi:10.1038/sj.emboj.7601798

18. Copley RR, Totrov M, Linnell J, Field S, Ragoussis J, Udalova IA. Functional conservation of Rel binding sites in drosophilid genomes. Genome Research. 2007;17: 1327–1335. doi:10.1101/gr.6490707

19. Tanji T, Yun E-Y, Ip YT. Heterodimers of NF-κB transcription factors DIF and Relish regulate antimicrobial peptide genes in Drosophila. Proceedings of the National Academy of Sciences. 2010;107: 14715–14720. doi:10.1073/pnas.1009473107

20. Ferrandon D, Jung AC, Criqui MC, Lemaitre B, Uttenweiler-Joseph S, Michaut L, et al. A drosomycin-GFP reporter transgene reveals a local immune response in Drosophila that is not dependent on the Toll pathway. EMBO Journal. 1998;17: 1217–1227. doi:10.1093/emboj/17.5.1217

21. Robinson SW, Herzyk P, Dow JAT, Leader DP. FlyAtlas: database of gene expression in the tissues of Drosophila melanogaster. Nucleic Acids Research. 2013;41: D744–D750. doi:10.1093/nar/gks1141

22. Levy F, Rabel D, Charlet M, Bulet P, Hoffmann JA, Ehret-Sabatier L. Peptidomic and proteomic analyses of the systemic immune response of Drosophila. Biochimie. 2004;86: 607–616. doi:10.1016/j.biochi.2004.07.007

23. Tzou P, Reichhart J-M, Lemaitre B. Constitutive expression of a single antimicrobial peptide can restore wild-type resistance to infection in immunodeficient Drosophila mutants. Proceedings of the National Academy of Sciences of the United States of America. 2002;99: 2152–2157. doi:10.1073/pnas.042411999

24. Wiegand I, Hilpert K, Hancock REW. Agar and broth dilution methods to determine the minimal inhibitory concentration (MIC) of antimicrobial substances. Nature Protocols. 2008. doi:10.1038/nprot.2007.521

25. Rabel D, Charlet M, Ehret-Sabatier L, Cavicchioli L, Cudic M, Otvos L, et al. Primary Structure and in Vitro Antibacterial Properties of the Drosophila melanogaster Attacin C Pro-domain. Journal of Biological Chemistry. 2004;279: 14853–14859. doi:10.1074/jbc.M313608200

26. Rahnamaeian M, Cytry ska M, Zdybicka-Barabas A, Dobslaff K, Wiesner J, Twyman RM, et al. Insect antimicrobial peptides show potentiating functional interactions against Gram-negative bacteria. Proceedings of the Royal Society B: Biological Sciences. 2015;282: 20150293–20150293. doi:10.1098/rspb.2015.0293

27. Kragol G, Lovas S, Varadi G, Condie BA, Hoffmann R, Otvos L. The antibacterial peptide pyrrhocoricin inhibits the ATPase actions of DnaK and prevents chaperone-assisted protein folding. Biochemistry. 2001;40: 3016–3026. doi:10.1021/bi002656a

28. Ryder E, Blows F, Ashburner M, Bautista-Llacer R, Coulson D, Drummond J, et al. The DrosDel collection: A set of P-element insertions for generating custom chromosomal aberrations in Drosophila melanogaster. Genetics. 2004;167: 797–813. doi:10.1534/genetics.104.026658

29. Ferreira ÁG, Naylor H, Esteves SS, Pais IS, Martins NE, Teixeira L. The Tolldorsal pathway is required for resistance to viral oral infection in Drosophila. PLoS Pathog. 2014;10: e1004507. doi:10.1371/journal.ppat.1004507

30. Levashina EA, Ohresser S, Bulet P, Reichhart J-M, Hetru C, Hoffmann JA. Metchnikowin, a Novel Immune-Inducible Proline-Rich Peptide from Drosophila with Antibacterial and Antifungal Properties. European Journal of Biochemistry. 1995;233: 694–700. doi:10.1111/j.1432-1033.1995.694_2.x

31. Fehlbaum P, Bulet P, Michaut L, Lagueux M, Broekaert WF, Hetru C, et al. Insect immunity: Septic injury of drosophila induces the synthesis of a potent antifungal peptide with sequence homology to plant antifungal peptides. Journal of Biological Chemistry. 1994;269: 33159–33163.

32. Zhang Y, Zhao J, Fang W, Zhang J, Luo Z, Zhang M, et al. Mitogen-Activated Protein Kinase hog1 in the Entomopathogenic Fungus Beauveria bassiana Regulates Environmental Stress Responses and Virulence to Insects. AEM. 2009;75: 3787–3795. doi:10.1128/AEM.01913-08

33. Dudzic JP, Hanson MA, Iatsenko I, Kondo S, Lemaitre B. More Than Black or White: Melanization and Toll Share Regulatory Serine Proteases in Drosophila. Cell Reports. 2019;27: 1050–1061.e3. doi:10.1016/j.celrep.2019.03.101

34. Binggeli O, Neyen C, Poidevin M, Lemaitre B. Prophenoloxidase Activation Is Required for Survival to Microbial Infections in Drosophila. PLoS Pathogens. 2014;10. doi:10.1371/journal.ppat.1004067

35. DeSimone SM, White K. The Drosophila erect wing gene, which is important for both neuronal and muscle development, encodes a protein which is similar to the sea urchin P3A2 DNA binding protein. Mol Cell Biol. 1993;13: 3641–3649. doi:10.1128/MCB.13.6.3641

36. Elya C, Lok TC, Spencer QE, McCausland H, Martinez CC, Eisen M. Robust manipulation of the behavior of Drosophila melanogaster by a fungal pathogen in the laboratory. eLife. 2018;7: e34414. doi:10.7554/eLife.34414

37. Imler J-L, Bulet P. Antimicrobial peptides in Drosophila: structures, activities and gene regulation. Chemical immunology and allergy. 2005;86: 1–21. doi:10.1159/000086648

38. Hedengren M, Borge K, Hultmark D. Expression and evolution of the Drosophila attacin/diptericin gene family. Biochemical and biophysical research communications. 2000;279: 574–81. doi:10.1006/bbrc.2000.3988

39. Khush RS, Lemaitre B. Genes that fight infection: what the Drosophila genome says about animal immunity. Trends Genet. 2000;16: 442–449. doi:10.1016/s0168-9525(00)02095-3

40. Goto A, Yano T, Terashima J, Iwashita S, Oshima Y, Kurata S. Cooperative regulation of the induction of the novel antibacterial Listericin by peptidoglycan recognition protein LE and the JAK-STAT pathway. J Biol Chem. 2010;285: 15731–15738. doi:10.1074/jbc.M109.082115

41. Barajas-azpeleta R, Wu J, Gill J, Welte R. Antimicrobial peptides modulate longterm memory. PLoS Genetics. 2018; 1–26. doi:10.1371/journal.pgen.1007440

42. Steffen H, Rieg S, Wiedemann I, Kalbacher H, Deeg M, Sahl H-G, et al. Naturally processed dermicidin-derived peptides do not permeabilize bacterial membranes and kill microorganisms irrespective of their charge. Antimicrob Agents Chemother. 2006;50: 2608–2620. doi:10.1128/AAC.00181-06

43. Zdybicka-Barabas A, Mak P, Klys A, Skrzypiec K, Mendyk E, Fiołka MJ, et al. Synergistic action of Galleria mellonella anionic peptide 2 and lysozyme against Gram-negative bacteria. Biochim Biophys Acta. 2012;1818: 2623–2635. doi:10.1016/j.bbamem.2012.06.008

44. Casteels-Josson K, Capaci T, Casteels P, Tempst P. Apidaecin multipeptide precursor structure: a putative mechanism for amplification of the insect antibacterial response. The EMBO journal. 1993;12: 1569–78. doi:10.1002/j.1460-2075.1993.tb05801.x

45. Hanson MA, Hamilton PT, Perlman SJ. Immune genes and divergent antimicrobial peptides in flies of the subgenus Drosophila. BMC evolutionary biology. 2016;16: 228. doi:10.1186/s12862-016-0805-y

46. Koganezawa M, Haba D, Matsuo T, Yamamoto D. The Shaping of Male Courtship Posture by Lateralized Gustatory Inputs to Male-Specific Interneurons. Current Biology. 2010;20: 1–8. doi:10.1016/j.cub.2009.11.038

47. Kurz CL, Charroux B, Chaduli D, Viallat-Lieutaud A, Royet J. Peptidoglycan sensing by octopaminergic neurons modulates Drosophila oviposition. Elife. 2017;6. doi:10.7554/eLife.21937

48. Lezi E, Zhou T, Koh S, Chuang M, Sharma R, Pujol N, et al. An Antimicrobial Peptide and Its Neuronal Receptor Regulate Dendrite Degeneration in Aging and Infection. Neuron. 2018;97: 125–138.e5. doi:10.1016/j.neuron.2017.12.001

49. Gramates LS, Marygold SJ, Dos Santos G, Urbano JM, Antonazzo G, Matthews BB, et al. FlyBase at 25: Looking to the future. Nucleic Acids Research. 2017;45: D663–D671. doi:10.1093/nar/gkw1016

50. Hill T, Koseva BS, Unckless RL. The genome of Drosophila innubila reveals lineage-specific patterns of selection in immune genes. Molecular Biology and Evolution. 2019. doi:10.1093/molbev/msz059

51. Chambers MC, Jacobson E, Khalil S, Lazzaro BP. Thorax injury lowers resistance to infection in Drosophila melanogaster. Infect Immun. 2014;82: 4380–4389. doi:10.1128/IAI.02415-14

52. Verleyen P, Baggerman G, D’Hertog W, Vierstraete E, Husson SJ, Schoofs L. Identification of new immune induced molecules in the haemolymph of Drosophila melanogaster by 2D-nanoLC MS/MS. Journal of Insect Physiology. 2006;52: 379–388. doi:10.1016/j.jinsphys.2005.12.007

53. Pfaffl MW. A new mathematical model for relative quantification in real-time RT-PCR. Nucleic Acids Res. 2001;29: e45. doi:10.1093/nar/29.9.e45

54. Tyler-Cross R, Schirch V. Effects of amino acid sequence, buffers, and ionic strength on the rate and mechanism of deamidation of asparagine residues in small peptides. J Biol Chem. 1991;266: 22549–22556.

