## Supplementary figures and images for "The Drosophila Baramicin polypeptide gene protects against fungal infection"

### FigureS1_Jan27-2021.pdf

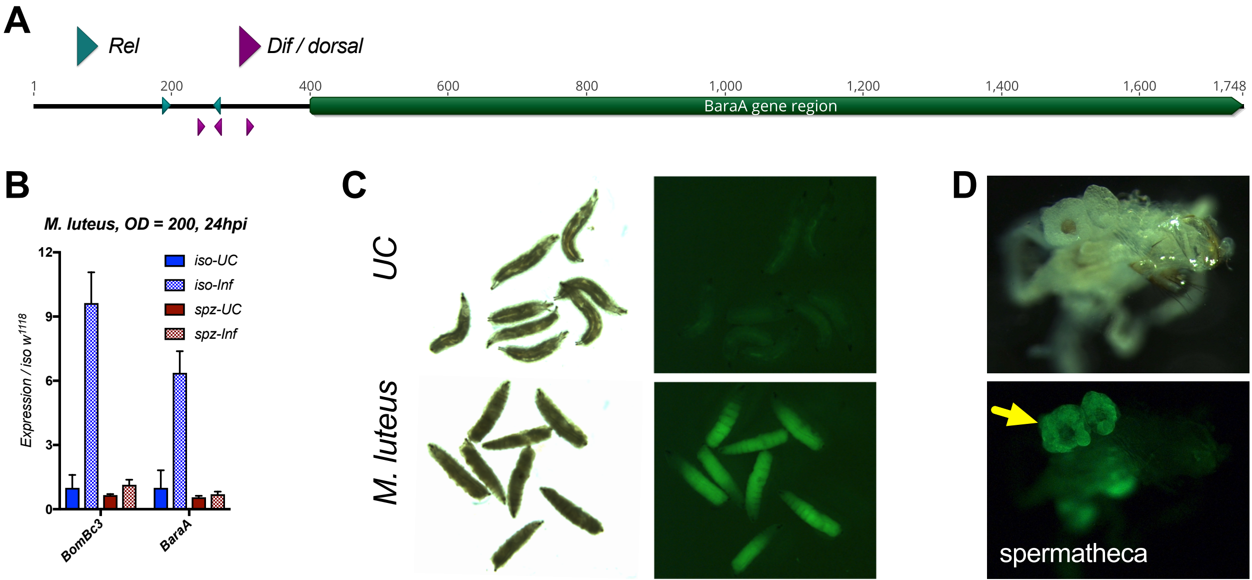

### FigureS2_BaraA_LCMScoverage.png

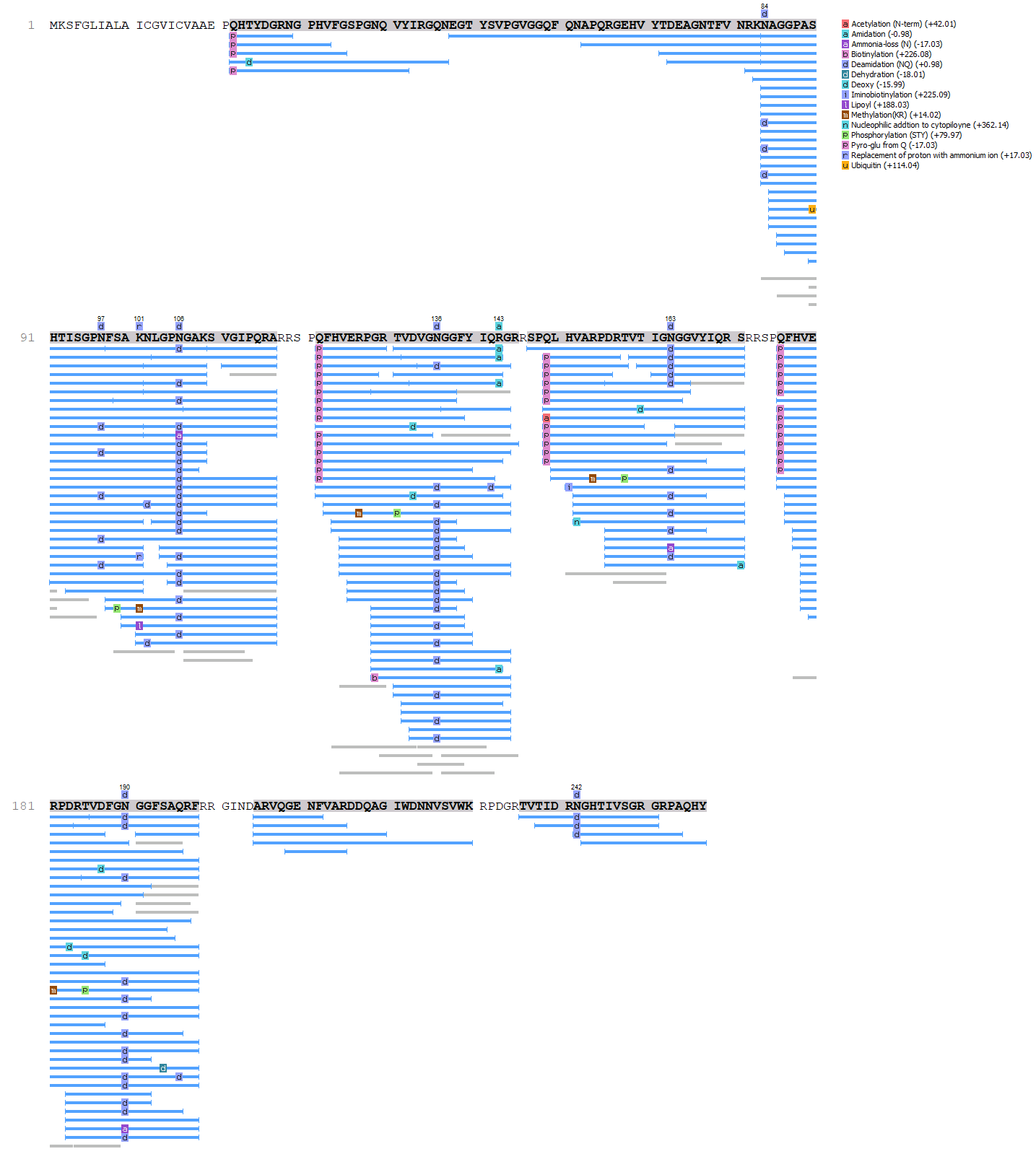

### FigureS3_IM22-IM10_Sept2020.pdf

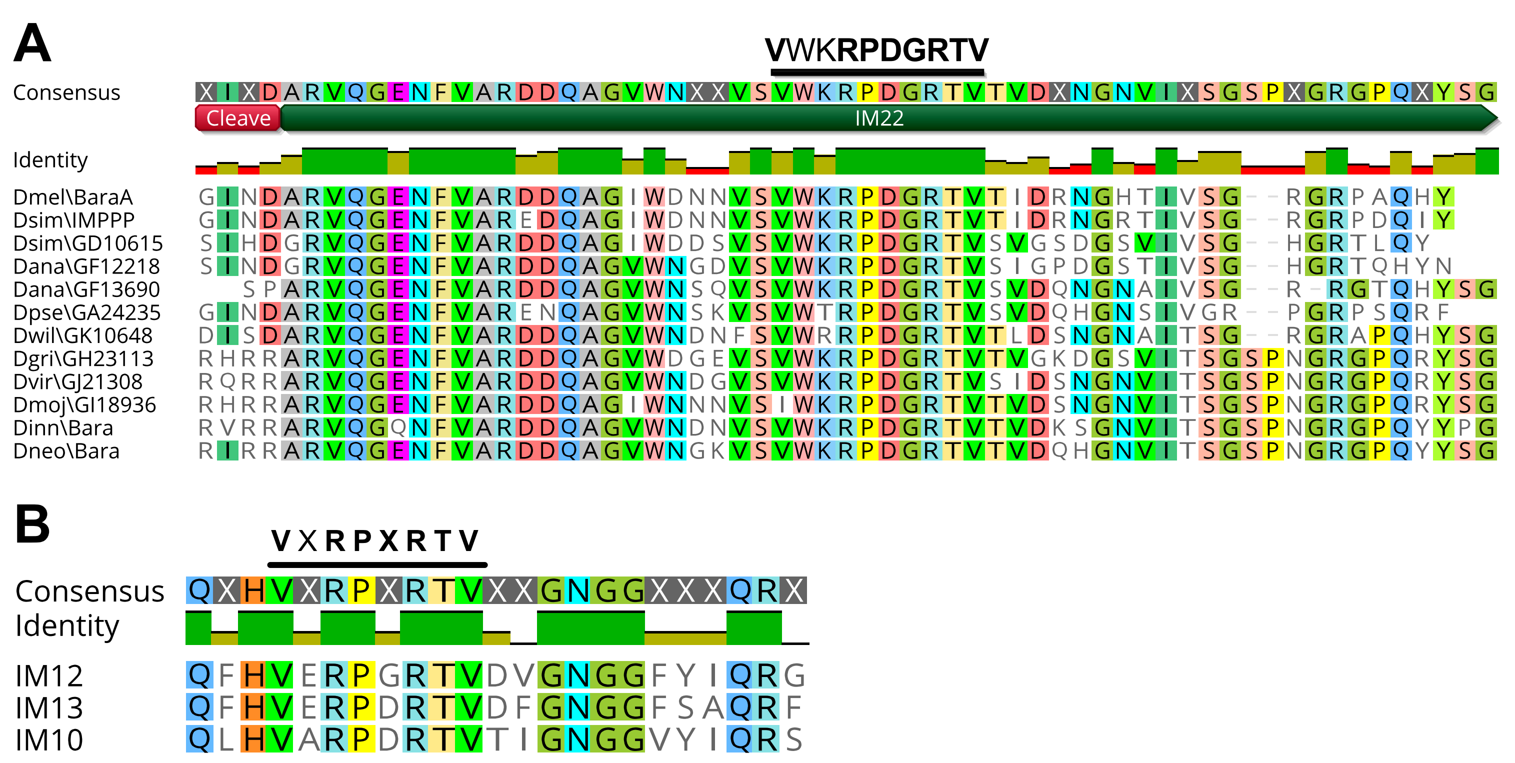

### FigureS4-ActBara_Relspz-Jan27-2021.pdf

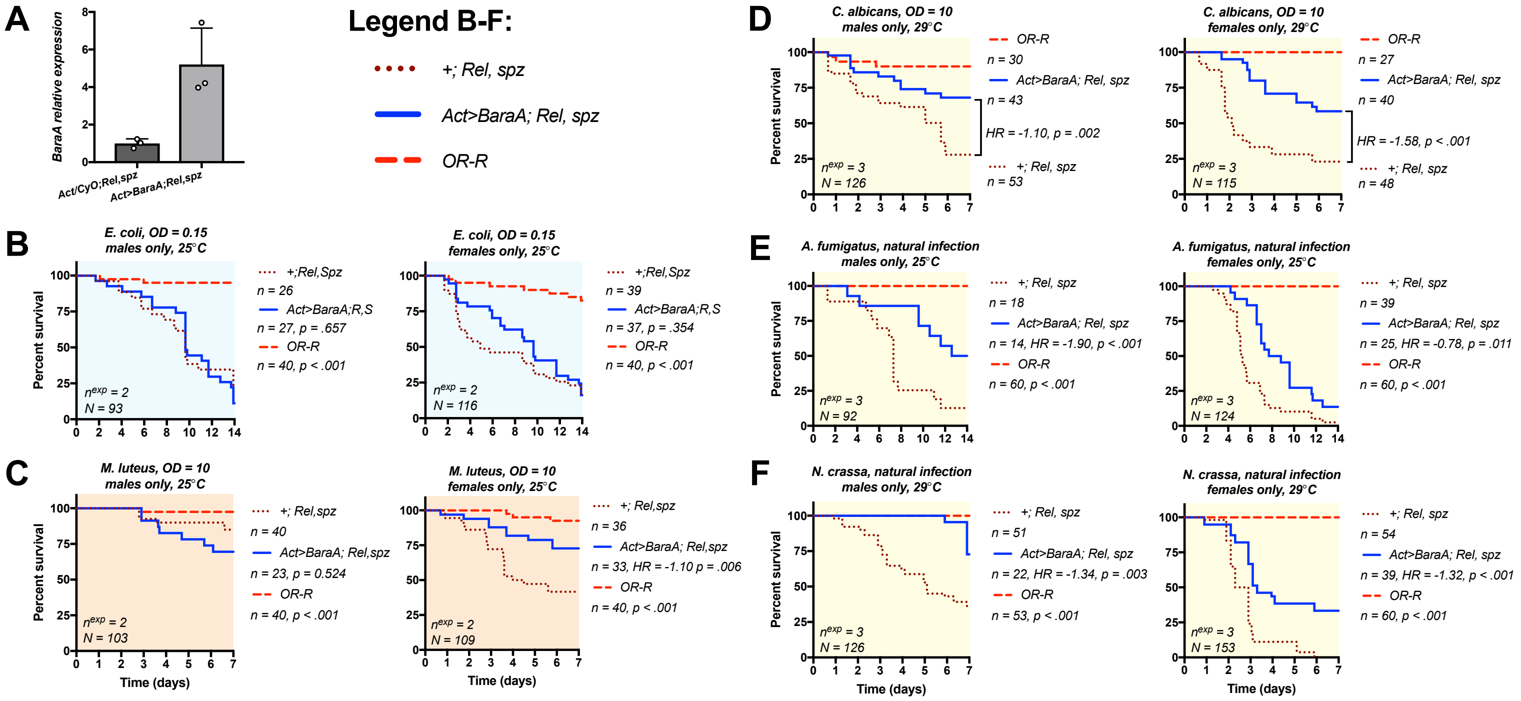

### FigureS5_qPCR-CI-Lifespan-Jan2021.pdf

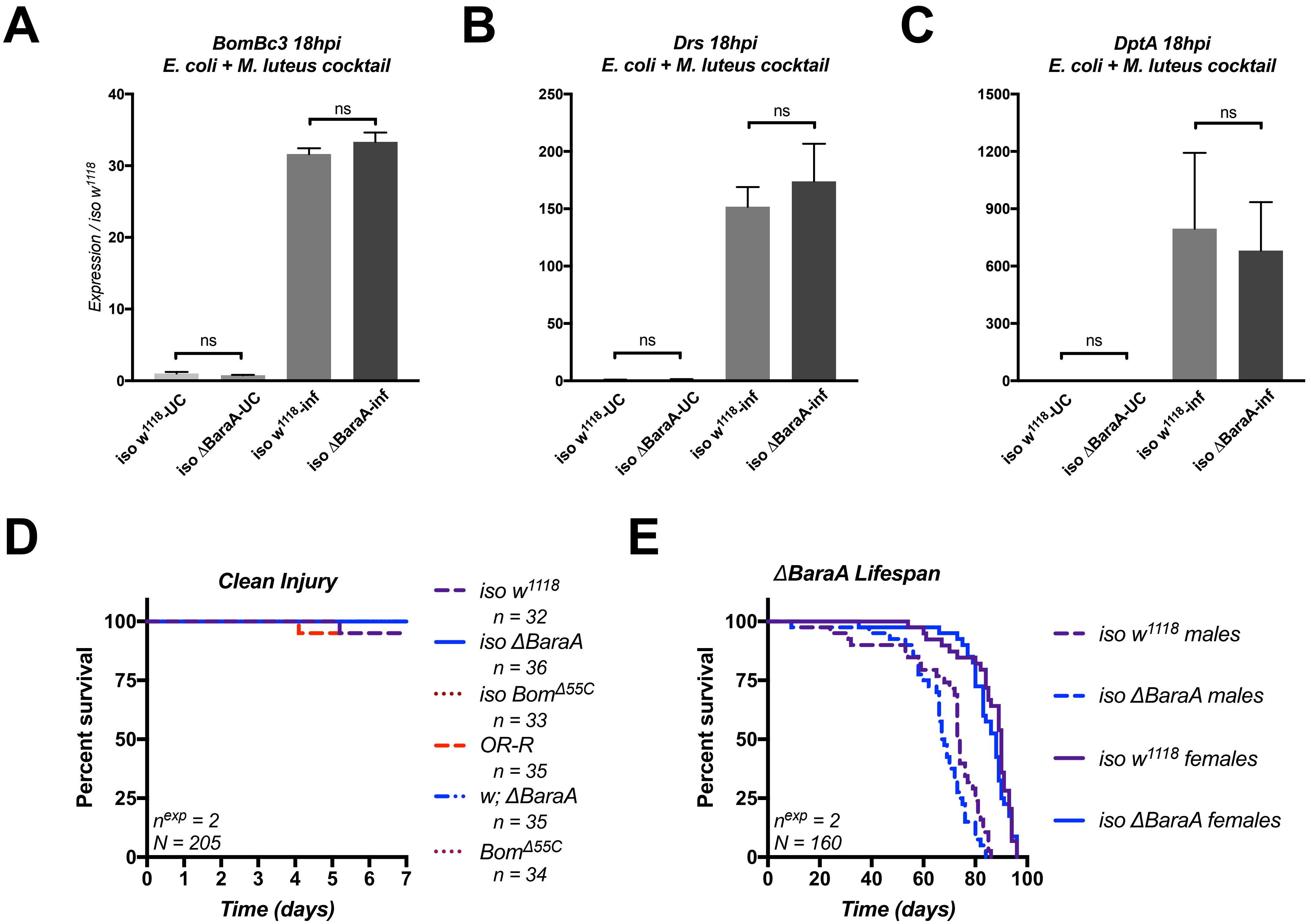

### FigureS6_Jan2021.pdf

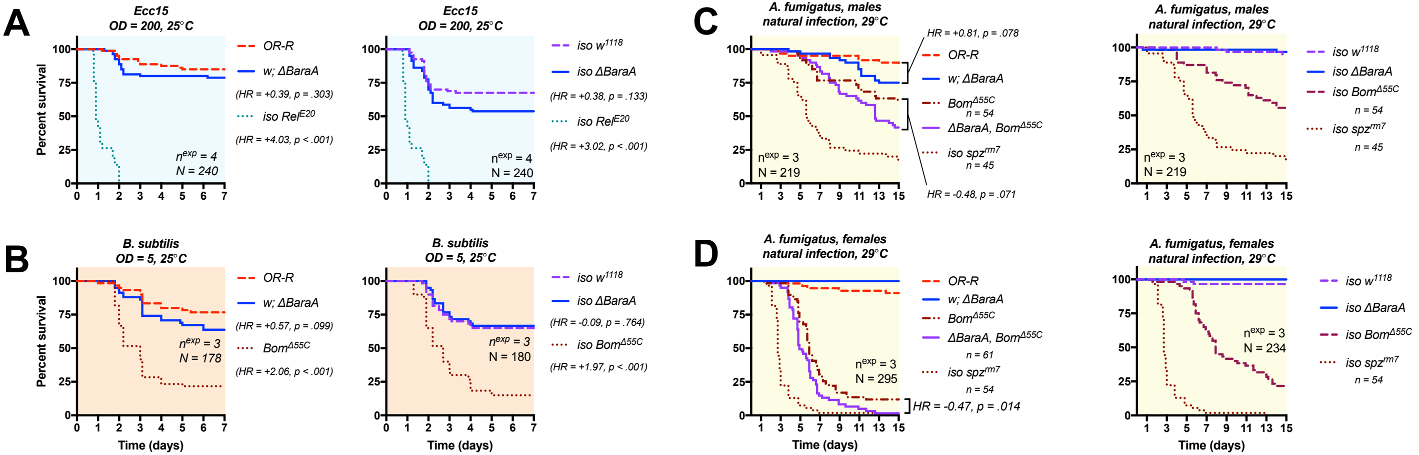

### FigureS7_Efae validation)Oct25_2020.pdf

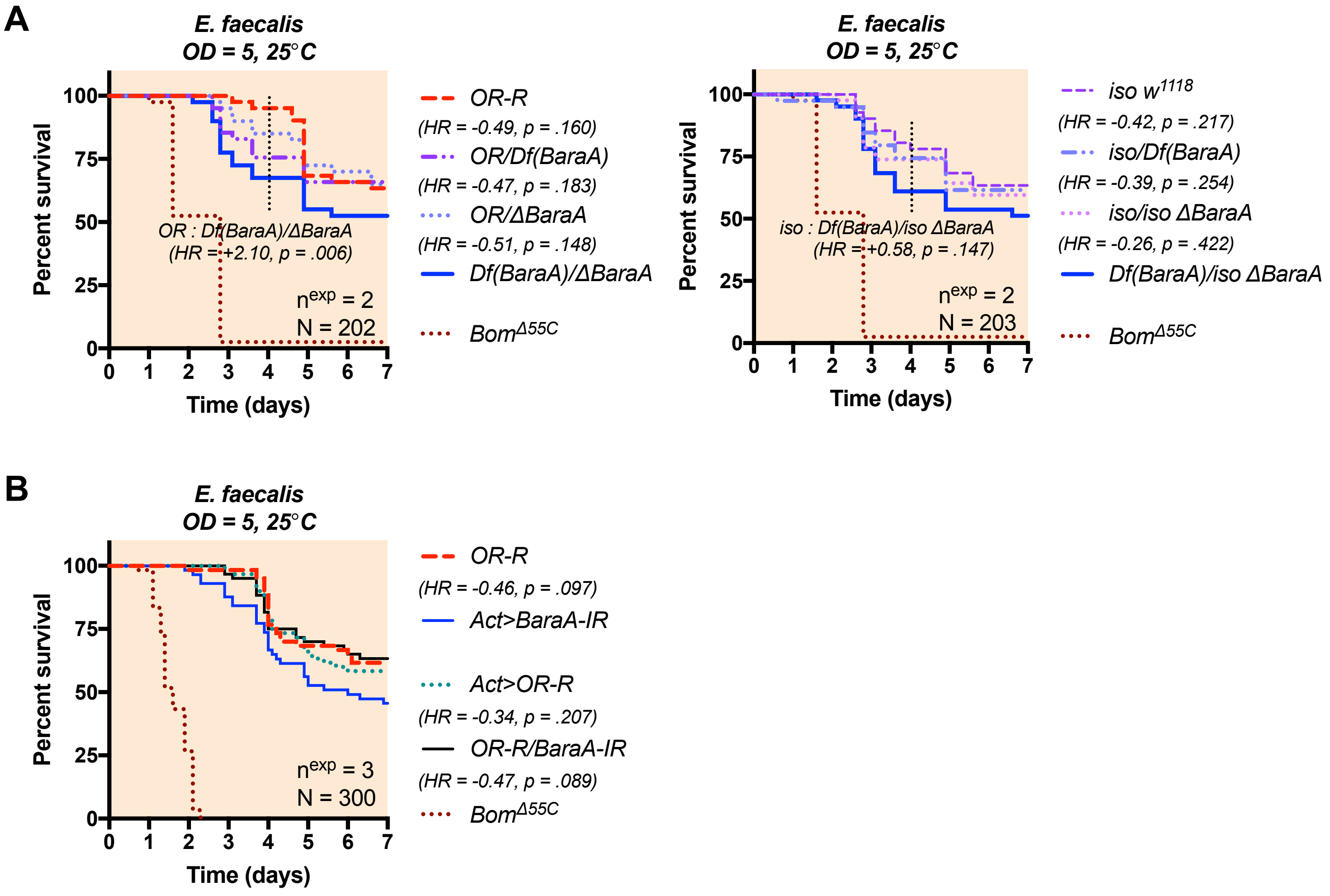

### FigureS8_Bbas validation-Jan2021.pdf

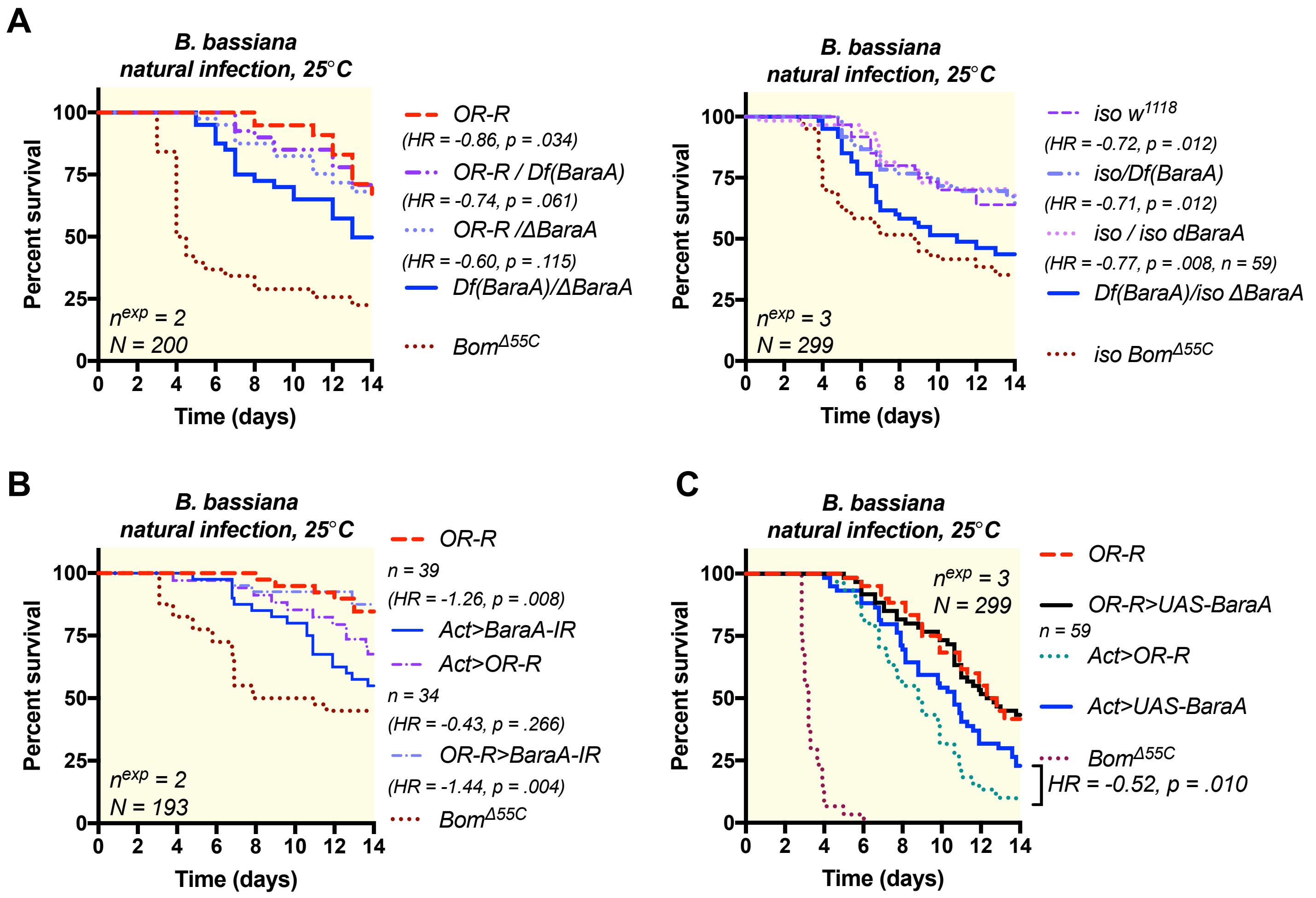

### FigureS9_erect wing-Jan2021.pdf

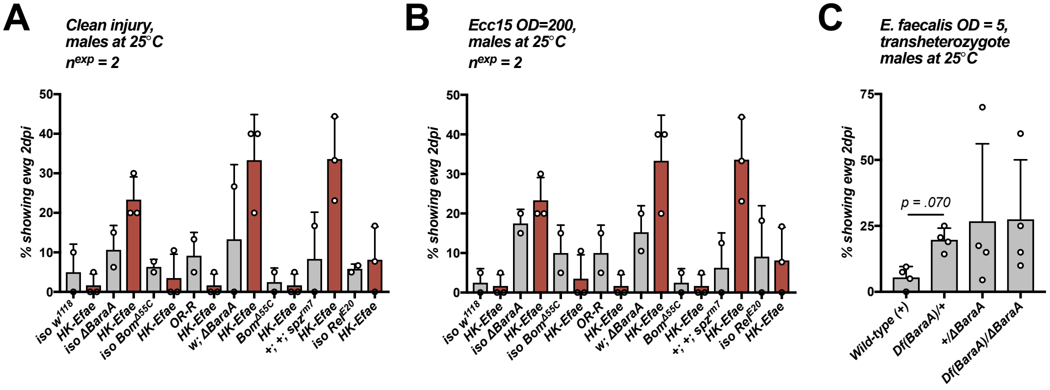

### TableS1_ewg_Jan-2021.png

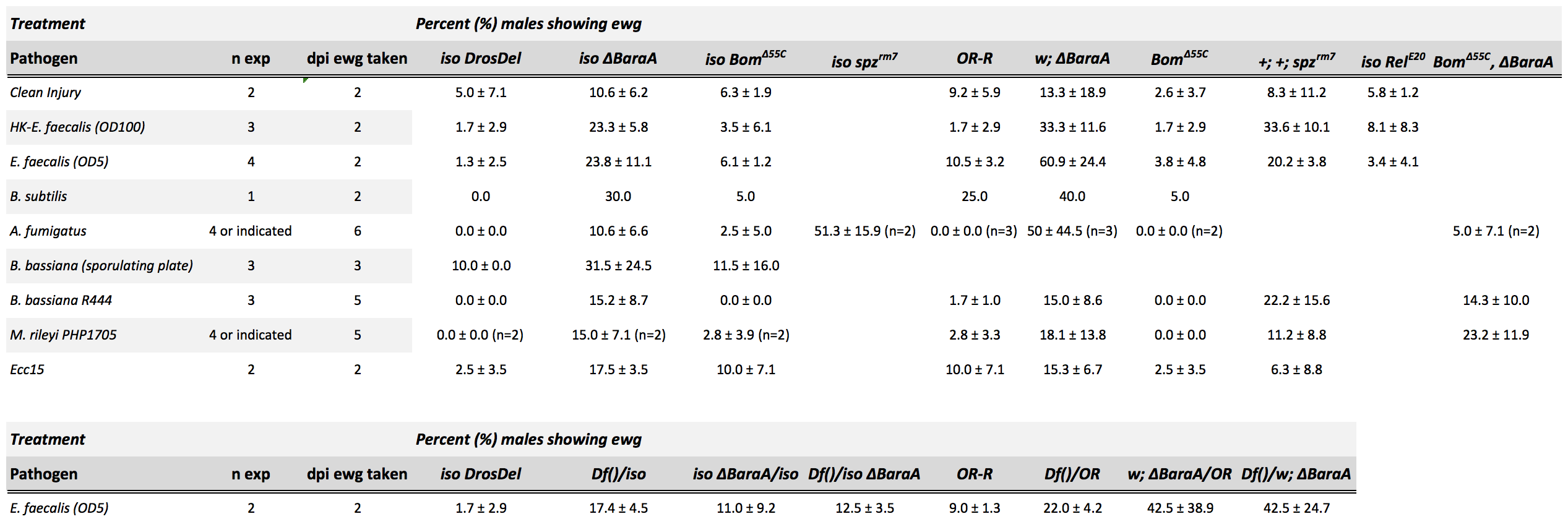
